# ChIMe-seq enables multimodal epigenetic profiling in single molecules and single cells

**DOI:** 10.64898/2026.11.17.767085

**Authors:** Joe Verity-Legg, Euan Joly-Smith, Roy Straver, Soufiane Mourragui, Stephen Clark, Vivek Bhardwaj, Marloes Blotenburg, Francesca Mattiroli, Sito Torres-Garcia, Wolf Reik, Jeroen de Ridder, Peter Zeller, Alexander van Oudenaarden

**Affiliations:** Hubrecht Institute-KNAW (Royal Netherlands Academy of Arts and Sciences), Oncode Institute, Utrecht, The Netherlands; University Medical Center Utrecht, Utrecht, The Netherlands; Oncode Institute, Utrecht, The Netherlands; Cambridge Institute of Science, Altos Labs, Cambridge, UK; Cancer Research UK Cambridge Institute, University of Cambridge; Cambridge, UK; Epigenetics Programme, The Babraham Institute, Cambridge, UK; Department of Physiology, Development and Neuroscience, University of Cambridge, Cambridge, UK

## Abstract

Recent advances in single-cell and single-molecule sequencing have enabled high-resolution characterization of epigenetic states. However, methods that jointly measure multiple layers of epigenetic regulation within the same molecules and cells remain limited. Here, we developed Chromatin Immuno-Methylation sequencing (ChIMe-seq), a multimodal approach that simultaneously profiles histone modifications, DNA methylation, and nucleosome positioning genome-wide in single molecules and single cells. In contrast to recent approaches that rely on enzymatic fragmentation, ChIMe-seq preserves long DNA molecules by using a methyltransferase to deposit exogenous methylation marks that report on proximal histone modifications. This enables integration of multiple epigenetic features within individual molecules and supports short-read, long-read, and single-cell sequencing. These multimodal measurements are integrated into a novel mathematical framework that captures the coupling between nucleosome positioning, histone modifications, and DNA methylation. Both single-molecule and single-cell analyses reveal that DNA methylation is linked to nucleosome density in a manner modulated by local histone modifications, whilst differences in nucleosome:DNA structures are observed depending on epigenetic context and cell-type. Together, these results establish ChIMe-seq as a framework to investigate multifaceted epigenetic coordination across scales, from single-nucleotide resolution to long-range chromatin organization in single molecules and single cells.

## INTRODUCTION

Advances in next-generation sequencing technologies have enabled systematic analysis of molecular layers that define cellular states and regulatory programs^1–3^. In particular, sequencing-based approaches have provided genome-wide measurements of transcription and its regulators. These measurements have revealed mechanisms of gene regulation across diverse biological systems.

More recently, single-cell and single-molecule technologies have further increased resolution, enabling the characterization of molecular heterogeneity within complex cell populations and the identification of regulatory states that are not apparent in bulk measurements^4,5^. These approaches allow direct observation of molecular states that exist within individual cells and along individual DNA molecules.

These developments have highlighted the interconnected nature of gene regulatory systems^6,7^. Cellular identity is controlled by multiple interacting layers of epigenetic regulators, including DNA methylation, histone modifications and nucleosome organization. To capture interactions between these layers, a growing number of multi-omic approaches have been developed that measure multiple molecular layers in parallel^8,9^. Despite this progress, methods that jointly profile histone modifications and DNA methylation remain limited. Existing approaches typically lack one or more key features, including: compatibility with long-read sequencing^10–12^, single-cell resolution^10,11,13–15^, high-resolution short-read sequencing^13–15^, ability to capture DNA methylation in the absence of histone modification target^10–12^, or the ability to capture nucleosome positioning^10,11^. As a result, it remains difficult to resolve how multiple epigenetic features are coordinated across scales ranging from single molecules to heterogeneous tissues. This limitation is particularly relevant given that epigenetic regulation is often context-dependent and varies across cell types, developmental stages, and disease states^16,17^.

Here, we developed Chromatin Immuno-Methylation sequencing (ChIMe-seq), a multimodal epigenetic profiling approach that simultaneously measures DNA methylation, histone modifications, and nucleosome positioning genome-wide in single molecules and single cells. In contrast to approaches that rely on enzymatic fragmentation of DNA, ChIMe-seq preserves long DNA molecules by using a methyltransferase-based strategy to deposit exogenous methylation marks that report on proximal histone modifications. This enables integration of multiple epigenetic features within individual DNA molecules while remaining compatible with short-read, long-read, and single-cell sequencing, allowing flexible experimental designs within a unified framework.

We demonstrate the accuracy of each ChIMe-seq modality in K562 cells and sensitively observe epigenetic state transitions, within individual DNA molecules, in long-reads. By developing a novel physical model of chromatin structure, we then quantitatively dissect the relationships between chromatin structure and epigenetic modifications with high resolution, revealing coordinated patterns between DNA methylation, histone modifications and nucleosome organization at the single-molecule level. Such relationships have, until now, not been possible to directly observe. Finally, we apply both ChIMe-seq and our physical model to mouse gastruloids to study these relationships in a heterogeneous population of single cells, revealing cell-type dependent relationships between epigenetic modifications and chromatin organization. Together, these results establish ChIMe-seq as a framework to investigate how epigenetic features are coordinated within chromatin across scales, from single-nucleotide resolution to long-range organization.

## RESULTS

### Development of a targetable GpC methyltransferase

ChIMe-seq builds upon previous work from our laboratories^18–20^ and others^21,22^ by using an antibody-directed methyltransferase to encode the presence of specific histone modifications directly into the associated DNA. This strategy enables the simultaneous measurement of histone modifications and endogenous DNA methylation on individual DNA molecules using a range of DNA methylation sequencing approaches (Fig. 1a).

**Figure 1.**
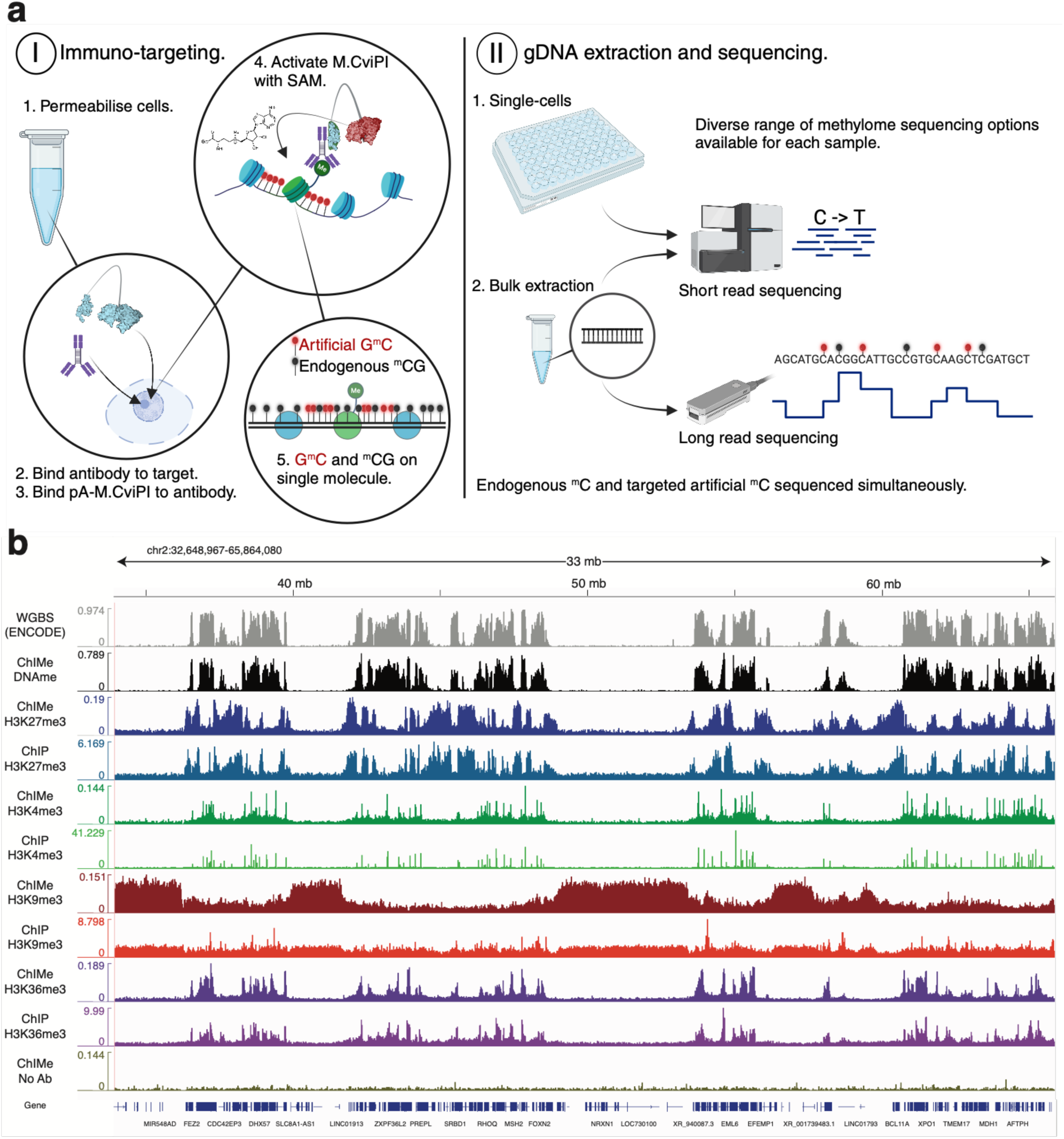
ChIMe-seq enables simultaneous recovery of multiple epigenetic features from a single experiment. **a.** Schematic overview of **(I)** the ChIMe-seq workflow and **(II)** downstream processing and sequencing strategies. **b.** Genome browser tracks showing ChIMe-seq profiles for multiple histone modification targets alongside corresponding ENCODE ChIP-seq datasets. ENCODE whole-genome bisulfite sequencing (WGBS) data are shown for comparison with the ChIMe-seq DNA methylation (DNAme) signal. The ChIMe-seq DNA methylation track is derived from the H3K27me3-targeted experiment. All tracks are displayed in 1-kb bins. y-axes indicate DNA methylation levels for methylation-based measurements and read counts for ChIP-seq datasets.

To achieve this, we engineered and purified a novel fusion protein consisting of Staphylococcal protein A and the GpC methyltransferase M.CviPI (pA-M.CviPI; Methods). Successful expression and purification of the 63-kDa protein were confirmed by gel electrophoresis (Supplementary Fig. 1a). The enzymatic activity of pA-M.CviPI was validated by incubation with unmethylated lambda phage DNA, followed by methylation-dependent restriction digestion using MspJI (Supplementary Fig. 1b). pA-M.CviPI exhibited activity comparable to that of commercially available M.CviPI, demonstrating that fusion to protein A does not compromise methyltransferase function.

In the ChIMe-seq workflow, a primary antibody is first bound to the histone modification of interest. pA-M.CviPI is subsequently recruited through the antibody-binding capacity of protein A. Upon activation, the tethered methyltransferase deposits GpC methylation near the targeted histone modification, thereby recording its genomic location. Because GpC methylation occurs in a sequence context distinct from the CpG methylation that predominates in mammalian genomes^1^, the two signals can be readily distinguished during downstream analysis. Furthermore, GpC methylation is preferentially deposited on DNA that is not protected by nucleosomes, as previously demonstrated^21,22^, providing a simultaneous readout of nucleosome positioning. As a result, histone modifications, endogenous DNA methylation, and nucleosome occupancy are all encoded within the same DNA molecule and can be measured in the same single cell.

### Validation of ChIMe-seq demonstrates its accuracy, sensitivity and reproducibility

To evaluate the accuracy of histone modification targeting by pA-M.CviPI while simultaneously measuring endogenous DNA methylation, we first performed bulk ChIMe-seq experiments in K562 cells targeting a range of histone modifications. ChIMe-seq GpC methylation profiles were compared with corresponding chromatin immunoprecipitation sequencing (ChIP-seq) datasets from the ENCODE consortium^23^, while DNA methylation (DNAme) profiles were compared with ENCODE whole-genome bisulfite sequencing (WGBS) data. Qualitatively, both ChIMe-seq modalities closely recapitulate the corresponding ENCODE reference datasets, whereas an antibody-free control exhibits minimal background signal, indicating low levels of non-specific methylation (Fig. 1b).

Quantitative comparison revealed strong positive correlations between ChIMe-seq and ENCODE datasets, with Pearson correlation coefficients of *r* = 0.65, 0.80, 0.84, 0.86, and 0.96 for H3K4me3, H3K27me3, H3K9me3, H3K36me3, and DNA methylation, respectively (Supplementary Fig. 2a–c). Consistent with their distinct chromatin distributions, GpC methylation profiles targeting heterochromatic marks showed little or negative correlation with profiles targeting euchromatic marks. DNA methylation displayed a strong positive correlation with H3K36me3 and anti-correlation with H3K27me3 and H3K9me3, in agreement with the established epigenetic landscape of K562 cells^16^ (Supplementary Fig. 2a,d).

To assess targeting specificity, we quantified the fraction of ChIMe-seq GpC methylation events and ChIP-seq reads that overlapped peaks called from ENCODE ChIP-seq datasets. ChIMe-seq exhibited specificity comparable to that of ChIP-seq when benchmarked against these reference peak sets (Supplementary Fig. 2e), whilst performing better or similarly than the reported performance of related methods (Supplementary Table 1). Sensitivity was assessed by measuring the fraction of GpCs within target peaks that were methylated. The observed values approached the theoretical maximum of approximately 25%, reflecting the fraction of genomic DNA that is accessible and not occupied by nucleosomes (Supplementary Fig. 2f).

To determine technical reproducibility, we compared independent ChIMe-seq replicates. Histone modification profiles (r = 0.97, Supplementary Fig. 2g) and DNA methylation profiles (r = 0.99, Supplementary Fig. 2h) showed high concordance between replicates. Importantly, DNA methylation measurements were highly consistent regardless of the histone modification being profiled, indicating that the endogenous DNA methylation readout is independent of the targeting strategy.

To evaluate protocol robustness and inter-laboratory reproducibility, ChIMe-seq was independently implemented in a second laboratory without prior training. Replicate experiments were performed in mouse 4T1 cells and benchmarked against matched CUT&Tag and WGBS datasets generated from the same cell culture. As observed in K562 cells, strong correlations were obtained between corresponding modalities (r = 0.62-0.99). Notably, ChIMe-seq identified H3K9me3 and H3K27me3 enriched regions unobserved by CUT&Tag, with CUT&Tag-based profiling of broad heterochromatic domains a known challenge (Supplementary Fig. 3).

Collectively, these results demonstrate that ChIMe-seq enables accurate, sensitive, and reproducible simultaneous measurement of histone modifications and DNA methylation. Furthermore, successful implementation in an independent laboratory highlights the robustness and transferability of the method.

### ChIMe-seq enables accurate and sensitive detection of DNA methylation and histone modifications across long single DNA molecules

Unlike many immuno-targeting methods, which identify targets through DNA fragmentation^18,24–27^, ChIMe-seq identifies targets by locally depositing methylation, preserving intact DNA molecules. As with other methyltransferase-based immuno-targeting approaches^13–15^, this enables long-read sequencing readouts (Fig. 1a), as well as detection of DNA methylation in the absence of histone modification targets.

Bisulfite conversion remains the gold-standard method for measuring cytosine methylation but substantially degrades DNA^28^, limiting long-read compatibility. Less destructive cytosine conversion methods exist^28–30^, supporting long-read sequencing^31,32^, although some enzymatic approaches show reduced performance in non-CpG contexts. Meanwhile, direct detection of DNA modifications by Oxford Nanopore Technologies (ONT) sequencing preserves native DNA and avoids amplification bias. We therefore evaluated the performance of ONT sequencing for ChIMe-seq by comparing matched Illumina and ONT datasets generated from the same sample.

Recent advances have substantially improved ONT-based DNA modification calling^33,34^, although model development has focused primarily on CpG methylation^35^. We found the publicly available GpC methylation model for R9 nanopore data (Remora, 5mC_all_context) accurately recapitulated genomic distributions of GpC methylation (*r* = 0.95), but markedly underestimated absolute methylation levels (Fig. 2a–c). We therefore trained a custom GpC methylation model (Methods), which increased detection sensitivity approximately sevenfold while maintaining the strong correlation with matched Illumina short-read measurements (*r* = 0.95) (Fig. 2c).

**Figure 2.**
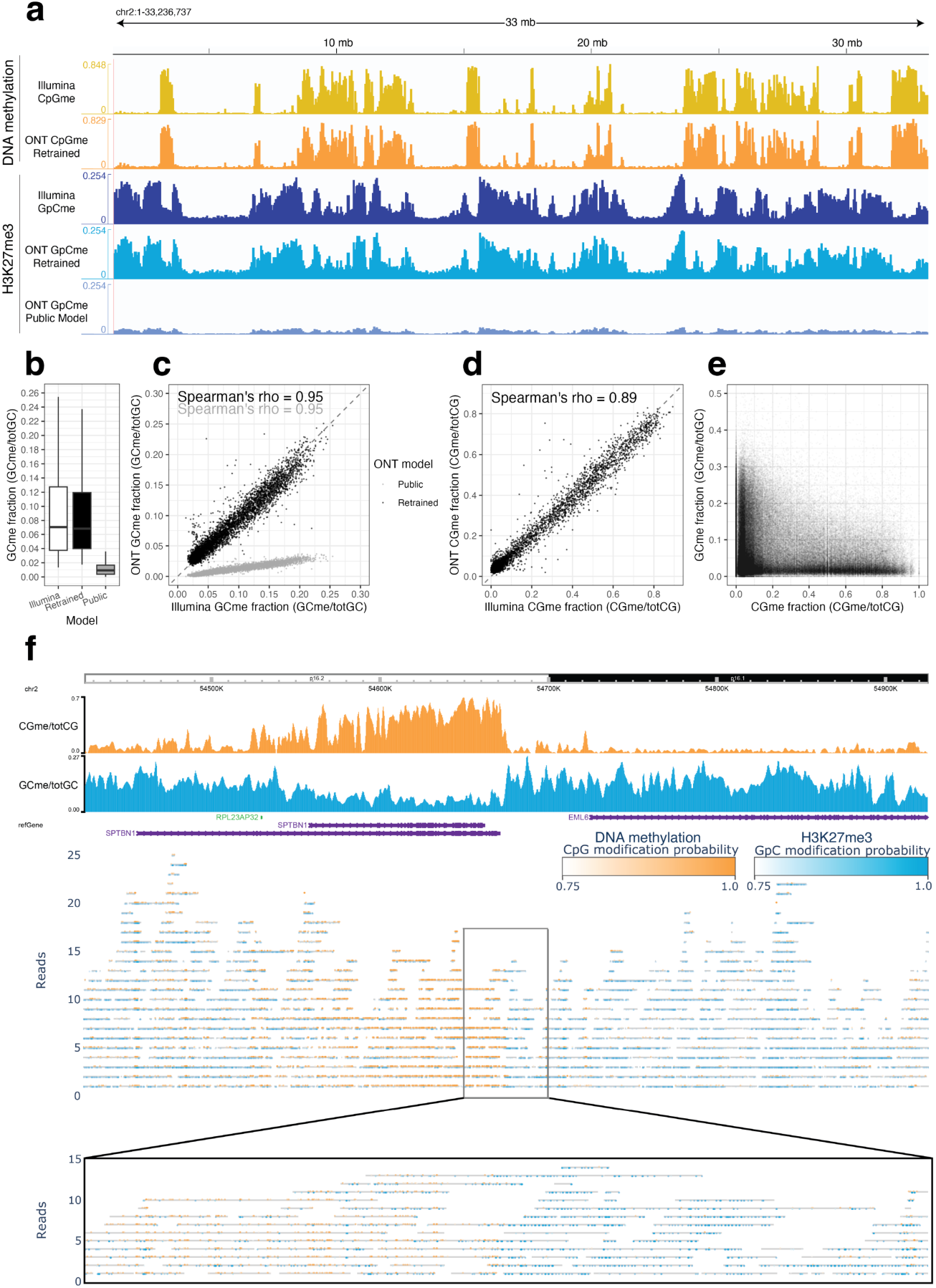
Non-destructive immuno-targeting enables long-read, single-molecule profiling of histone modifications and DNA methylation. **a.** Genome browser tracks of a H3K27me3-targeted ChIMe-seq sample analysed by whole-genome bisulfite sequencing (Illumina) or native DNA methylation detection using Oxford Nanopore Technologies (ONT). **b.** Distribution of GpC methylation levels across 50-kb bins on chromosome 2 for Illumina sequencing and ONT sequencing using the public and retrained GpC methylation models. **c.** Scatter plot comparing Illumina and ONT GpC methylation measurements across 50-kb bins on chromosome 2. Grey points indicate comparisons using the public ONT model, whereas black points indicate the retrained model. **d.** Scatter plot comparing Illumina and ONT CpG methylation measurements across 50-kb bins on chromosome 2. **e.** Scatter plot of the mean GpC and CpG methylation levels for individual sequencing reads. Each point represents a single DNA molecule. **f.** Single-molecule visualization of CpG and GpC methylation across a 500-kb region of chromosome 2. The upper panel shows aggregate CpG methylation (orange) and GpC methylation (blue) tracks in the WashU Epigenome Browser. Individual read alignments are displayed below in grey. Coloured markers denote methylated sites with a posterior probability ≥0.75 (orange, CpG; blue, GpC), with colour intensity proportional to methylation probability. The inset shows a higher-resolution view of the central 50-kb region, illustrating simultaneous detection of CpG and GpC methylation on individual DNA molecules.

Assessing CpG methylation calling in ChIMe-seq data, the state-of-the-art R9 CpG methylation model (Remora) consistently overcalled methylated cytosines in some sequence contexts, relative to Illumina sequencing (Supplementary Fig. 4b-c). For both v1 and v2 of the public CpG model overcalling was most pronounced in regions also containing GpC methylation, particularly in NHCGC sequence contexts where GpC and CpG motifs occur in close proximity. We therefore hypothesized that neighbouring GpC methylation may elicit false-positive CpG methylation calls in ONT data. To address this, we fine-tuned a Remora model using lambda phage genome artificially methylated in GpC and / or CpG contexts as training data. Thus, the model was taught to recognise the correct methylation state of CpG sites proximal to GpC methylation.

Following fine-tuning, CpG methylation became underestimated in HCCGN and CACGN contexts (Remora v2 retrained, Supplementary Fig. 4b-c), possibly in part due to CpG methylation contamination of the GpC methylation only training dataset, due to non-GpC activity of the M.CviPI enzyme (Supplementary Fig. 4d). These contexts were excluded from downstream analyses. The resulting CpG methylation model showed a very strong correlation with matched Illumina short-read measurements (r = 0.89, Fig. 2d).

Using our optimized GpC and CpG methylation models, we simultaneously profiled H3K27me3 and endogenous DNA methylation on individual DNA molecules. ChIMe-seq generated long sequencing reads with a median length of 3,684 bp and an N50 of 15,604 bp. At the single-molecule level, H3K27me3-associated GpC methylation and endogenous CpG methylation were found to be largely mutually exclusive, consistent with their known genomic distributions (Fig. 2e).

Finally, we visualized individual GpC and CpG methylation events on single DNA molecules spanning an epigenomic domain boundary (Fig. 2f). The improved sensitivity of our custom models enabled clear delineation of domain boundaries on individual molecules, demonstrating accurate and simultaneous detection of multiple epigenetic features within the same long DNA fragment.

### ChIMe-seq reveals nucleosome organization across epigenetic states

Nucleosomes are organized into continuous, spatially coordinated arrays fundamental to chromatin structure and function^36–39^. However, although CpG methylation, histone modifications and nucleosome organization are proposed to be interconnected, these relationships have not been directly observed in a cellular context, limiting our understanding of such relationships. Because pA-M.CviPI preferentially methylates nucleosome-free linker DNA, ChIMe-seq captures patterns of nucleosome organization around targeted histone modifications. This enables quantitative characterization of chromatin architecture across multiple epigenetic modalities, in long, single DNA molecules.

To investigate nucleosome organization within discrete epigenetic contexts, long ChIMe-seq reads were segmented *in silico* into 1-kb fragments. Fragments from experiments targeting four different histone modifications were stratified by their CpG methylation levels, enabling comparisons across distinct epigenetic states. Within each fragment, the methylation state of each GpC was recorded along with its distance from every other methylated GpC. Aggregating these measurements across fragments within a CpG methylation / histone modification group yielded pairwise conditional methylation probabilities, describing the probability that a GpC is methylated as a function of its genomic distance from another methylated GpC. The resulting curves exhibited characteristic damped oscillations (Supplementary Fig. 5), reflecting periodic alternation of nucleosome-protected and linker DNA, consistent with previous observations^22,40^. Notably, these oscillations varied according to histone modification and CpG methylation level, indicating that nucleosome organization differs between epigenetic states (Supplementary Fig. 5). In contrast, shuffling methylation statuses of individual GpCs while preserving their genomic positions abolished oscillatory structures, confirming they reflect chromatin organization rather than sequence-derived biases (Supplementary Fig. 6).

To interpret these patterns, and decipher the epigenetic context specific differences in chromatin structure they represent, we developed a physical model of nucleosome organization (Fig. 3a). In this model, nucleosome footprints and linker DNA form a repeating chain. Footprint and linker lengths are modelled as independent, gamma-distributed, random variables. The mean nucleosome footprint length is fixed at 147 bp, the canonical length of DNA wrapped around histone octamers. Within modified domains, GpCs are methylated by pA-M.CviPI with probabilities *P_nuc_* and *P_linker_*, depending on whether they fall within nucleosome-protected or linker DNA. GpCs outside modified domains are assumed to be unmethylated. Finally, to account for the finite extent of histone-modified domains, we introduce a boundary probability: the probability per base of leaving the modified domain when moving away from the methylated GpC used to anchor the conditional probability.

**Figure 3.**
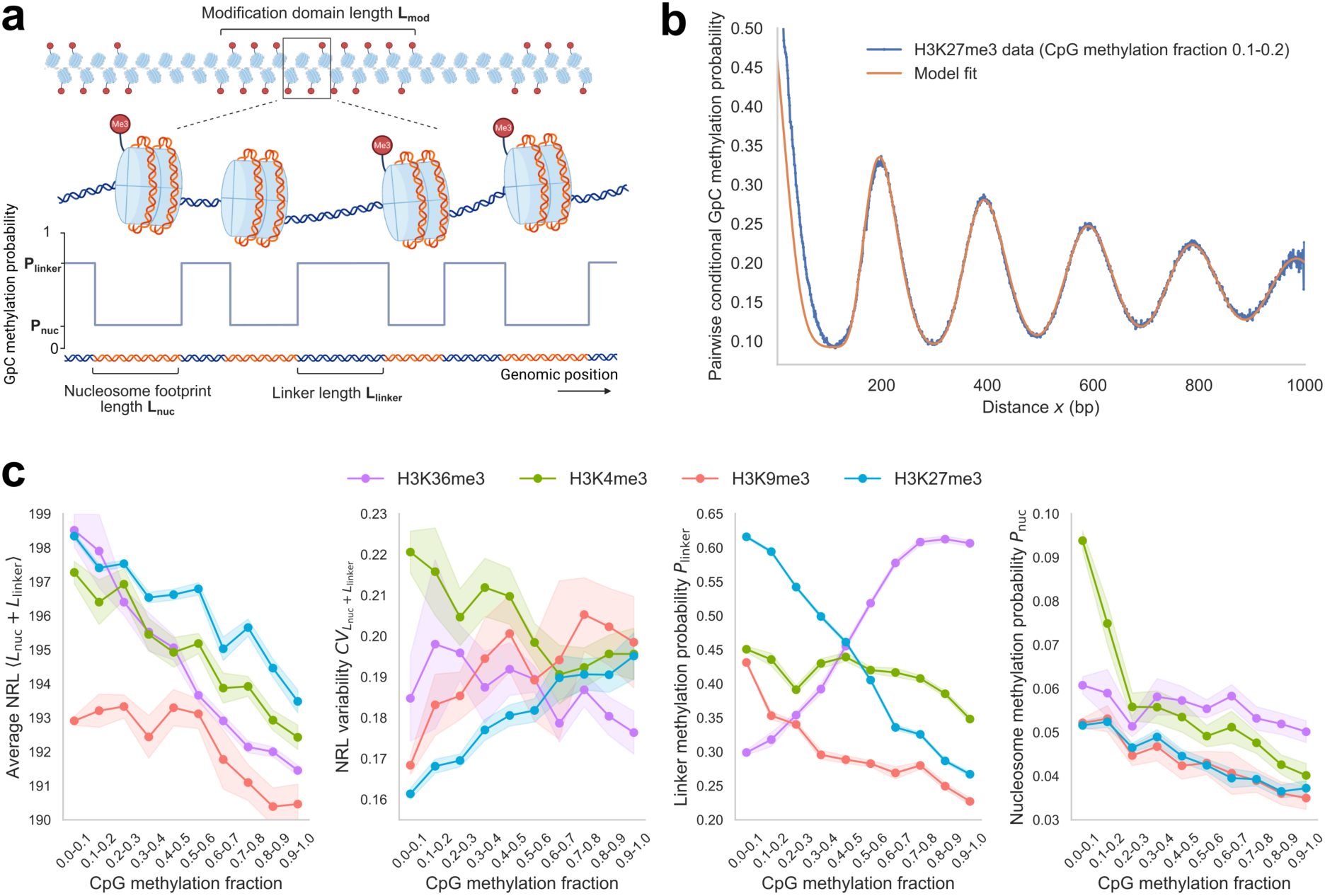
Nucleosome organization quantified from single-molecule GpC methylation. **a.** Schematic of the nucleosome-linker model. To account for the finite extent of histone-modified domains, a boundary probability describes the probability per base pair of leaving the modified domain when moving away from the methylated GpC used to anchor the conditional probability. A region of length L_mod_ is decomposed into nucleosome footprints (orange) and linker DNA (blue). Footprint and linker lengths L_nuc_ and L_linker_ are gamma-distributed. The average nucleosome footprint length is fixed at 147 bp, corresponding to canonical DNA wrapping around the histone octamer. Within modified regions, GpCs are methylated with probabilities P_nuc_ and P_linker_ depending on whether they fall within nucleosome-protected or linker DNA. GpCs outside modified domains are unmethylated. **b.** Pairwise conditional methylation probability as a function of genomic distance x, computed across H3K27me3 reads with CpG methylation fraction 0.1-0.2 (blue curve). The oscillatory structure reflects the periodic alternation of nucleosome-protected and linker DNA. The orange curve shows the analytical model prediction fit by least-squares optimization. **c.** Inferred model parameters as a function of CpG methylation fraction across histone modifications. Error bars indicate 95% confidence intervals estimated from the Jacobian-based covariance matrix of the least-squares fit. Angle brackets denote the mean, whereas CV denotes the coefficient of variation of the indicated quantity. <*L*_nuc_ + *L*_linker_> denotes the average nucleosome repeat length, whereas CV_Lnuc+Llinker_ denotes its coefficient of variation.

We derived a closed-form analytical expression from the model for the pairwise conditional methylation probability (Methods). Fitting this expression to the observed oscillatory profiles using least-squares optimization allowed direct inference of model parameters (Fig. 3b). In contrast, many existing methods rely on hidden Markov models to segment nucleosome and linker regions. These models implicitly assume exponential length distributions^41^ which is inconsistent with known nucleosome organization^42–44^. Our approach is conceptually related to hidden semi-Markov models^41^ and provides a general framework for non-exponential state durations. We leveraged the analytical expression of pairwise methylation probabilities for efficient parameter inference.

Applying this framework across fragment groups of varying CpG methylation fraction and histone modification revealed systematic variation in inferred parameters, predicting fine-grain structural features of chromatin, even with sparse GpC motif data (Fig. 3c, Supplementary Fig. 7).

First, we find the average nucleosome repeat length (NRL) broadly decreases with increasing CpG methylation across histone modifications (Fig. 3c, Average NRL). However, specifically in H3K9me3-marked chromatin, NRLs are both notably shorter and relatively unchanged with low CpG methylation before decreasing at higher methylation levels. This indicates broadly increased nucleosome density (through shorter linkers) with increasing CpG methylation, revealing genome-wide relationships between DNA methylation and chromatin organization.

Variability in nucleosome spacing is the primary driver of oscillation decay, with domain boundaries providing an additional contribution. In K562 cells, heterochromatin becomes increasingly disordered with increasing CpG methylation, whereas euchromatin shows the opposite trend, becoming more regularly organized (Fig. 3c, NRL variability). Domain organization also differs between chromatin states: at low CpG methylation, H3K4me3 shows the smallest domains, with highest boundary density (λ), whereas H3K27me3 shows the largest domains (Supplementary Fig. 7, l).

Linker GpC methylation probability (*P_linker_*) also varies across chromatin states. Because pA-M.CviPI is recruited to the targeted histone modification, higher *P_linker_* is expected in regions containing a greater density of modified nucleosomes. H3K36me3 *P_linker_* increases with CpG methylation, whereas H3K27me3 and H3K9me3 *P_linker_* decreases, consistent with their respective enrichment across CpG methylation states (Fig. 3c, *P_linker_*).

Finally, nucleosome GpC methylation probability (P*_nuc_*) reports the likelihood that GpCs within nucleosome-protected DNA are methylated by pA-M.CviPI. Subtle but consistent differences are observed across epigenetic states (Fig. 3c, P_nuc_). P*_nuc_* is highest for H3K4me3 at low CpG methylation, whereas H3K27me3 and H3K9me3 show lower P*_nuc_* than euchromatic modifications. Across all histone modifications, P*_nuc_* decreases with increasing CpG methylation, suggesting reduced accessibility of nucleosome-protected DNA to pA-M.CviPI.

In summary, we find that CpG methylation is consistently related to shorter NRLs and reduced accessibility of nucleosomal DNA to pA-M.CviPI across epigenetic states. Euchromatic nucleosomal DNA appears more accessible than heterochromatic, particularly at putative active gene promoters, whilst the relationship between the regularity of nucleosome spacing and CpG methylation levels differs between euchromatin and heterochromatin. Together, these results demonstrate that ChIMe-seq, and our mathematical framework, enable quantitative characterization of nucleosome organization across diverse epigenetic states.

### ChIMe-seq accurately identifies cell-lineages from single cells in mouse gastruloids

To demonstrate the ability of ChIMe-seq to resolve epigenetic states at single-cell resolution, we applied it to a heterogeneous system, an *in vitro* model of mouse gastrulation. We first showed effective antibody targeting in gastruloids via flow cytometry, finding 99.47% of cells were stained (Supplementary Fig. 8a). Next, we processed 864 single ChIMe-seq cells (targeting H3K27me3) with scBS-seq^19,20^ to yield single-cell methylation data. Across all cells, CpG methylation is depleted at transcription start sites (TSSs), and GpC methylation is enriched across H3K27me3 peaks called from reference data^45^ (Supplementary Fig. 8b-c), confirming broad accuracy of both modalities. After quality filtering, retaining 76% of cells, a median of 297,407 CpGs and 2,679,963 GpCs were observed across 660 cells (Supplementary Fig. 8d-f). Finally, comparison of pseudobulked single-cells and bulk sequencing of the same ChIMe-seq preparation showed similar methylation rates across contexts (Supplementary Fig. 8g).

CpG and GpC methylation calls were analysed independently generating modality-specific single-cell count matrices (Methods). A joint CpG–GpC UMAP embedding resolved four distinct cell clusters (Fig. 4a). Cluster separation was reduced when modalities were analysed separately with at least one population poorly defined, demonstrating that multi-omic ChIMe-seq outputs can improve the resolution of cellular heterogeneity in single-cell experiments.

**Figure 4.**
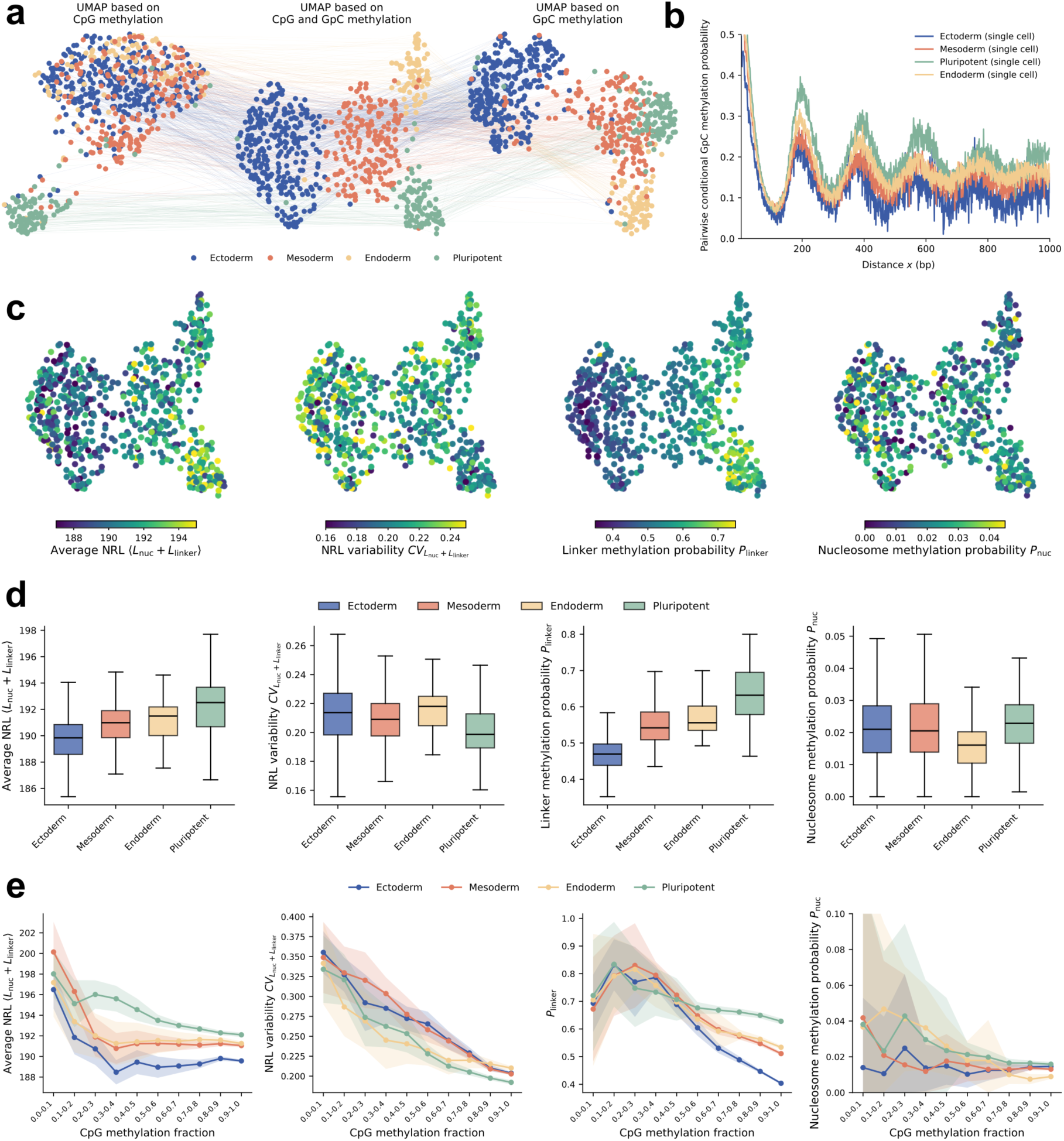
Single-cell integration of DNA methylation, H3K27me3 and chromatin structure in day 5 mouse gastruloids. **a.** Joint analysis of CpG and GpC methylation profiles from ChIMe-seq single cells. UMAP embeddings generated from CpG-only, GpC-only and integrated CpG–GpC datasets are shown, with corresponding cells linked across representations. Four distinct cell clusters are resolved in the integrated embedding. **b.** Representative conditional GpC methylation probability curves from individual cells, showing oscillatory patterns consistent with nucleosome organization within H3K27me3-marked chromatin. **c.** Integrated CpG–GpC UMAP embedding coloured by fitted model parameters, including average NRL, NRL variability, linker methylation probability (P_linker_), and nucleosome methylation probability (P_nuc_). **d.** Box plots showing the distribution of single-cell fitted parameter values across gastruloid lineages. Box plots show the median (center line) and interquartile range (box), with whiskers extending to 1.5x the interquartile range. **e.** Nucleosome organization as a function of DNA methylation state. Lines and points represent mean parameter estimates across 100 bootstrap refits, generated by resampling cells and repeating CpG methylation stratification, conditional GpC methylation probability calculation and model fitting. Shaded regions indicate ±1 s.d. across bootstrap refits.

To assign lineage identities to the four clusters, our results were compared with a reference T-ChIC dataset^45^, which simultaneously measured RNA and H3K27me3 in single cells. ChIMe-seq H3K27me3 profiles showed strong lineage-specific correlations with T-ChIC H3K27me3 profiles (Supplementary Fig. 9a). Further, promoter-proximal H3K27me3 and CpG methylation signals showed lineage-specific anti-correlations with T-ChIC RNA expression from the associated genes (Supplementary Fig. 9b-c), with the exception of endoderm CpG methylation. Thus, ChIMe-seq epigenetic signals may be linked to transcriptional states in single cells. Most importantly, the lineage of the four ChIMe-seq populations was assignable using only scRNA-seq reference data, and the expected cell populations of the cellular model were recovered. Interestingly, the proportions of cells assigned to each lineage for day 5 gastruloid ChIMe-seq, closely resembles those from RNA based assignments of day 6 gastruloid T-ChIC – suggesting that epigenetic changes broadly precede transcriptional changes in this system (Supplementary Fig. 9d).

### ChIMe-seq enables quantitative assessment of chromatin structure, across epigenetic environments, in single-cells

Having established lineage assignments, we sought to investigate nucleosome organization in single-cells through application of our physical model of chromatin structure. For single cells, we accounted for a lack of phasing of our short-read data through introduction of a synchronization parameter, *k*. *k* is the probability that a genomic position is in a linker on one chromosome given that the corresponding position is in a linker on the other chromosome. Lower synchronization reduces the oscillatory amplitude and was included in the single-cell model (Methods). The distribution of GpC methylation again displayed characteristic, oscillatory nucleosome patterns in randomly selected single cells from each lineage (Fig. 4b). Subsequently, fitting our physical model provided quantitative predictions for structural chromatin features, across H3K27me3 DNA in single cells (Fig. 4c). This revealed clear trends in lineage-dependent variations in chromatin structure (Fig. 4d).

Firstly, pluripotent cells possessed the highest average NRL for H3K27me3 chromatin, with ectoderm cells the lowest. Pluripotent cells also displayed lower NRL variability across H3K27me3 heterochromatin than other cell types. Opposing trends in P*_linker_* and domain boundary density (λ) were found, with this relationship observable across individual cells (Supplementary Fig. 10b), suggesting cells with larger H3K27me3 domains also have more densely modified domains. Finally, changes in P*_nuc_* are observed across lineages. Absolute P*_nuc_* differences are small, but vary across single cells. On average, P*_nuc_* is higher in pluripotent cells and lower in endoderm cells, suggesting differences in H3K27me3-modified nucleosome structures across lineages.

Finally, nucleosome organization was analysed as a function of DNA methylation state within each lineage (Fig. 4e, Supplementary Fig 10c). Due to the sparsity of single-cell methylation data, CpG methylation calls were aggregated across cells, within each lineage, in 1 kb bins, before stratification of bins by CpG methylation fraction. The single-cell model was fitted, yielding nucleosome organization parameters as a function of DNA methylation state, within each lineage. As in our K562 long-read data, NRL decreases with increasing CpG methylation, across lineages. Despite different epigenetic modifications being linked to differential NRLs within an individual cell type (Fig. 3c), differences in NRL between lineages remain when measurements are restricted to H3K27me3 DNA with similar CpG methylation levels. In contrast to K562s, H3K27me3 NRL variability decreases with increasing CpG methylation, across all gastruloid lineages. Although no clear relationship between H3K27me3 P*_nuc_* and CpG methylation was observed in single cells, unlike in our K562 dataset, P*_nuc_*showed large errors arising from finite cell sampling, preventing reliable estimation of this parameter. However, as expected, H3K27me3 P*_linker_* decreases with increasing CpG methylation, as was also observed for K562 cells (Fig. 3c).

In summary, we find that in pluripotent cells H3K27me3 chromatin is less densely populated by nucleosomes, but that H3K27me3 domains are more densely modified, when compared to the three germ layers. Further, pluripotent H3K27me3 chromatin has the most regularly arranged nucleosome arrays as well as nucleosome core DNA that is the most accessible. Conversely, ectoderm H3K27me3 chromatin is most densely populated with nucleosomes, but have H3K27me3 domains that are the least densely modified. We also find that nucleosome core DNA in endoderm H3K27me3 chromatin is the least accessible. Finally, our single-cell data supports our previous observation that increasing CpG methylation is associated with more densely arranged nucleosomes.

Our results show the capabilities of ChIMe-seq, combined with our novel mathematical framework to deliver unprecedented resolution for dissecting chromatin structure, in single cells.

## DISCUSSION

We present ChIMe-seq, a multi-omic sequencing method for simultaneous profiling of histone modifications and DNA methylation (DNAme). ChIMe-seq provides specific, sensitive, reproducible, and robust measurements of both epigenetic modalities. Long-read ChIMe-seq enables sensitive detection of histone modifications and reveals epigenetic domain boundaries on individual DNA molecules. At single-cell resolution, the unique combination of genome-wide histone modification and DNAme profiles distinguishes individual cell lineages in mouse gastruloids, with cell type resolution improved when compared with a unimodal readout.

To leverage the nucleosome-footprinting capability of ChIMe-seq, we developed a mathematical framework for the quantitative dissection of chromatin structure. Applying this framework to both long-read and single-cell data, we examine the interplay between nucleosome organization, histone modifications, and DNAme, providing an unprecedented view of how these three chromatin features are coordinated. We extend recent observations linking CpG methylation and nucleosome spacing^46^ showing that increasing CpG methylation is related to a dose-dependent shortening of NRLs within single chromatin fibres. CpG methylation is reported to intrinsically alter higher-order genome conformations^47^ whilst changes in linker length, of the size we observe between fully methylated and unmethylated DNA, are also proposed to produce significant inter-nucleosomal conformational shifts^48^. ChIMe-seq extends these observations by resolving the relationship between DNAme and nucleosome spacing quantitatively and, most importantly, within defined histone-modification contexts. NRL decreases approximately linearly with increasing DNAme in H3K4me3-, H3K36me3-, and H3K27me3-marked chromatin, whereas H3K9me3-marked chromatin displays a distinct relationship, with NRL remaining relatively constant at low DNAme before decreasing at higher methylation levels. These H3K27me3 results are reproduced across lineages, in single cells. Thus, CpG methylation might offer a general means to progressively tune chromatin structure, both locally and globally, across cell-types. We also observe subtle shifts in the probability of nucleosomal GpCs being methylated by pA-M.CviPI across distinct epigenetic contexts. We interpret higher nucleosomal GpC methylation as reflecting increased accessibility of nucleosome-associated DNA to the M.CviPI methyltransferase. These differences may therefore report chromatin context-dependent variation in nucleosome structure or dynamics that expose otherwise protected DNA to enzymatic modification, in line with recent results^49^. Consistent with this interpretation, P*_nuc_* is elevated in H3K4me3-marked chromatin at low DNA methylation levels. This may reflect the increased structural dynamics of nucleosomes at active promoters, where fragile nucleosomes have been described and proposed to facilitate rapid transcriptional responses^50,51^. In vitro experiments and simulations have suggested that CpG methylation can alter DNA wrapping around nucleosomes^52,53^, although both the direction and magnitude of these effects remain debated^54^. Rather than directly reporting DNA wrapping, ChIMe-seq provides a functional measure of the resulting accessibility of nucleosome-associated DNA to enzymatic interaction in native chromatin. Similarly, simulations have suggested that histone-tail modifications can modulate histone–DNA and nucleosome–effector interactions^55^. Our results extend these structural predictions to individual nucleosomes in cells, revealing that the biochemical accessibility of nucleosomal DNA varies systematically with both DNA methylation and histone-modification state. In general, we find repressive epigenetic modifications (DNAme, H3K9me3, H3K27me3) are related to reduced biochemical accessibility of nucleosomal DNA. Together, these findings support a model in which epigenetic modifications not only mark distinct chromatin states, but are associated with differences in the physical accessibility of DNA within individual nucleosomes, providing a potential additional layer of chromatin regulation.

In summary, our findings support a growing body of evidence that epigenetic features act in concert to tune dynamic local and higher-order chromatin architecture, with potential consequences for genome regulation. By combining ChIMe-seq with a quantitative framework for analysing methylation-footprinting data, we directly examine the interplay between histone modifications, DNAme, nucleosome structure, and nucleosome array organization at single-molecule and single-cell resolution. Our results reveal previously inaccessible relationships between these chromatin features and demonstrate how their integration can provide a quantitative view of chromatin organization across multiple structural scales. More broadly, our mathematical framework is applicable to existing and future methylation-footprinting methods^13–15,22,56–68^, providing a general approach for extracting quantitative information about nucleosome organization and accessibility from this rapidly expanding class of assays.

## Methods

### Cloning of pA-M.CviPI

pK19pA-M.CviPI was cloned by replacing micrococcal nuclease of pK19pA-MN (Plasmid #86973 – Addgene) with the sequence for M.CviPI taken from Genbank, AF062394. A construct containing: part of protein A (from an AfeI site downstream), a flexible linker containing a His tag and M.CviPI followed by a downstream BamHI site was ordered from IDT. pK19pA-MN and the M.CviPI construct were digested with BamHI and AfeI, the purified vector was dephosphorylated and the new insert ligated into the vector. The resulting plasmid pK19pA-M.CviPI was cloned into chemically competent DH5α bacteria for amplification, using kanamycin for selection. Sequence integrity was validated by sanger sequencing. For protein production chemically competent E. coli strain C3013 was transformed with pK19pA-M.CviPI. The full protein sequence is available in Supplementary Note.

### pA-M.CviPI production

400mL pre-culture was grown overnight in LB broth with 1:1000 kanamycin and 1% (w/w) glucose at 37°C whilst shaking (200rpm). Pre-cultures were then centrifuged (3220g, 10mins, 4°C). Supernatant was discarded and cells re-suspended in 200mL fresh LB broth before transfer to 3.2L LB broth with 1:1000 kanamycin. Cultures were grown to OD_600_ 0.8-1.0 at 37°C whilst shaking (200rpm). Protein expression was induced with IPTG (Isopropyl beta-D-1-thiogalactopyranoside) at a final concentration of 2mM and culture incubated for 2-3 hours at 37°C whilst shaking (200rpm). Cells were centrifuged (3220g, 10mins, 4°C) and pellet re-suspended in 160mL lysis buffer (10mM Tris-HCl pH 7.5, 150mM NaCl, 5mM MgCl_2_, 5mM CaCl_2_, 12µL (3000 U) Pierce Universal Nuclease (Thermo Scentific 88700), 5mM DTT and 4 Roche cOmplete EDTA-free Protease Inhibitor Cocktail tablets (11873580001), 10% glycerol (v/v)). Lysozyme (Sigma, L4919) was added to 50µg/mL and cell suspension was incubated on ice for 10 mins. Cells were lysed on ice in 40mL aliquots using a Branson Sonifier SFX 150 (with a 1/4“ microtip) (2 pulses, 40s on, 3min off, 70% amplitude). Lysed cells were centrifuged (13000g, 25mins, 4°C). Lysate was filtered through a 22µm filter. 3mL of Chelating Sepharose Fast Flow resin (Cytiva, 17-0575-02) was mixed with 3mL 100mM Ammonium Ni (II) Sulfate Hexahydrate charging buffer (Sigma, 574988), mixed well by inversion and placed on ice for 3mins. Resin was washed 3x with 40mL, 4°C miliQ water and 2x with Ni Buffer A (10mM Tris-HCl pH 7.5, 150mM NaCl, 10mM imidazole, 1mM TCEP, 10% glycerol (v/v)) (all centrifugations: 1100g, 4mins, 4°C). Resin was mixed with filtered cell lysate and incubated for 1hr at 4°C with rotation. Resin-lysate mixture was applied to a gravity column (Bio-Rad, 9704652), washed with 160mL Ni Buffer A and eluted in 13mL Ni Buffer B (10mM Tris-HCl pH 7.5, 150mM NaCl, 300mM imidazole, 1mM TCEP, 10% glycerol (v/v)). Eluent was loaded onto a 1mL HiTrap Heparin HP column (Cytiva, 17040601), pre-equilibrated with 10 column volumes Heparin Buffer B (10mM Tris-HCl pH 7.5, 1M NaCl, 1mM TCEP, 10% glycerol (v/v)) and 10 column volumes Heparin Buffer A (10mM Tris-HCl pH 7.5, 150mM NaCl, 1mM TCEP, 10% glycerol (v/v)). The column was washed with 15 column volumes Heparin Buffer A before pA-M.CviPI was eluted using a linear gradient of 0-100% Heparin Buffer B across 15 column volumes. 0.5 mL fractions were collected. Fractions with A280 absorbance were analysed by denaturing gel electrophoresis using Bolt 4-12%, Bis-Tris 1.0mm Mini Protein Gels (Invitrogen, NW04127BOX) in NuPage MES SDS Running Buffer (Invitrogen, NP000202) and with a Precision Plus Protein Dual Color Standards ladder (Bio-Rad, 1610374). Samples had SDS loading buffer added to final concentration of 50mM Tris-HCl (pH 6.8) 100mM DTT, 2% SDS 0.1% bromophenol blue, 10% glycerol, were heated to 98°C for 3mins and briefly centrifuged. Gel was run at 200V for 30mins, stained using InstantBlue Coomassie Protein Stain (Abcam, ab119211) and imaged using an Amersham ImageQuant 800. Product containing fractions were measured with the Qubit Protein Assay Kit and made up to final storage conditions of 1µg/mL pA-M.CviPI in 15mM Tris-HCl pH 7.5, 200mM NaCl, 1mM DTT, 200 µg/mL BSA, 0.1mM EDTA, 50% glycerol (v/v) and snap-frozen in liquid N_2_ for storage at −80°C. Enzymatic activity of each fraction was tested using the *in vitro* activity assay.

### pA-M.CviPI activity assay

1µg Non-methylated Lambda Phage gDNA (Sigma, D3654-5UN) in 20µL with 1x GC reaction buffer (NEB, B0227S), 3.2mM SAM (NEB, B9003S) and 1µL enzyme (either 4U M.CviPI (NEB, M0227) or 1ng pA-M.CviPI) or H_2_O was incubated for 4hrs at 37°C. 10μL of each sample was transferred to 10μL MspJI digestion mix: 1X rCutSmart Buffer (NEB, B6004S), 2x Enzyme Activator (NEB, S0538S) and 5U MspJI (NEB, R0661). The MspJI reaction was heated at 37°C for 2hrs and 65°C for 20mins. Samples were analysed using Agilent Bioanalyzer HS DNA kit (5067-4626).

### Cell culture

K562 cells (ATCC® CCL-243™) were grown in RPMI 1640 Medium GlutaMAX™, supplemented with 5% FCS, Pen-Strep and non-essential amino acids. The mouse mammary tumour cell line 4T1 (ATCC Cat# CRL-2539, RRID:CVCL_0125) was cultured in high-glucose DMEM (41966-029, Gibco), supplemented with 10% heat-inactivated fetal bovine serum (A5209501, Gibco) and 50 U/mL penicillin-streptomycin (15140122, Gibco). The cell line was STR profiled to confirm identity and tested negative for mycoplasma. After harvesting cells were washed 3 times with room temperature PBS before continuing with the ChIMe-seq protocol.

### Cell preparation: fixation

All buffers are made fresh for each experiment, rolled at 4°C for 30mins before use and kept at 4°C or on ice throughout protocol unless otherwise stated. Steps were performed as for Cell preparation: fixation in sortChIC^13^ but with ChIC wash buffer as: 20mM HEPES pH 7.5 (Gibco, 15630080), 150mM NaCl, 4mM EDTA, 0.459mM spermidine, 0.05% Tween20, 1 Roche cOmplete EDTA-free Protease Inhibitor Cocktail tablet (11873580001) per 50 ml buffer. Cells were centrifuged at 600g, 4°C, 3mins for this and all further centrifugations in the protocol, unless otherwise stated.

### pA-M.CviPI targeting

Cells were pelleted and re-suspended to 1.75M cells / mL in ChIC wash buffer. 400µL per desired sample was aliquoted into Eppendorf 0.5mL Protein LoBind tubes (Eppendorf, 0030108094) and antibody was added to the following final dilutions: H3K27me3 – 1:100 - H3K27me3 Rabbit mAB (C36B11) (CellSignaling, #9733), H3K9me3 – 1:200 - H3K9me3 Recombinant Rabbit Monoclonal Antibody (RM389) (Invitrogen, MA5-33395), H3K4me3 – 1:400 - H3K4me3 Monocloncal Antibody (G.532.8) (Invitrogen, MA5-11199), H3K36me3 – 1:2000 - H3K36me3 Recombinant Rabbit Monocolonal Antibody (RM155) (Invitrogen, MABEE640). Samples were mixed by pipetting ten times and incubated overnight at 4°C on a roller. Samples were pelleted and washed 1x with ChIMe wash buffer (50mM Tris-HCl pH 7.5, 150mM NaCl, 10mM DTT, 4mM EDTA, 0.459mM spermidine, 0.005% Tween20, 1 Roche cOmplete EDTA-free Protease Inhibitor Cocktail tablet (11873580001) per 50 ml buffer. After wash samples were centrifuged again and pellet re-suspended in 500µL ChIMe wash buffer. 2.5ng pA-M.CviPI was added to each sample and mixed by pipetting. Samples were incubated at 4°C for 1hr with rolling. Cells were washed 2x with ChIMe wash buffer and finally re-suspended in ChIMe wash buffer. Samples were incubated at 35°C for 25mins, washed 2x with 35°C ChIMe wash buffer and 35°C centrifugation before finally being re-suspended in room temperature ChIMe reaction buffer (50mM Tris-HCl pH 7.5, 50mM NaCl, 10mM DTT, 4mM EDTA, 0.459mM spermidine, 0.005% Tween20, 1 Roche cOmplete EDTA-free Protease Inhibitor Cocktail tablet (11873580001) per 50 ml buffer). Fresh SAM (S-adenosyl methionine) was added to a final concentration of 320µM and mixed well. Samples were incubated at 25°C for 4hrs. Reactions were stopped by pelleting and washing 1x at 4°C with stop buffer (50mM Tris-HCl pH 7.5, 300mM NaCl, 4mM EDTA, 0.05% Tween20, 1 Roche cOmplete EDTA-free Protease Inhibitor Cocktail tablet (11873580001) per 50 ml buffer). Samples were spun down and re-suspended based upon their downstream usage.

### Bulk short-read ChIMe-seq

gDNA was extracted with Qiagen DNeasy Blood & Tissue Kit (69504) following manufacturer’s guidelines, quantified with Qubit dsDNA HS Assay Kit (10616763) and processed with Zymo Research Pico Methyl-Seq Library Prep Kit (D5456) using 100ng input and following the manufacturer’s guidelines. Abundance and quality of the final library were assessed by Qubit dsDNA HS Assay Kit (10616763) and Agilent Bioanalyzer HS DNA kit (5067-4626) and sequenced paired-end 2×150bp using an Illumina NextSeq500.

### CUT&Tag

CUT&Tag was performed on 100,000 cells per condition and replicate following a published protocol^69^. Cells were detached with 0.05% Trypsin for 5 minutes at 37°C, centrifuged at 600 g for 3 mins at room temperature, washed with 1 volume of PBS, and resuspended in ½ volume of ice-cold NE1 buffer (20 mM HEPES-KOH pH 7.5, 10 mM KCl, 0.5 mM spermidine, 0.1% Triton X-100, 20% glycerol, 1x EDTA-free Complete Protease Inhibitor). After a 10-minute incubation on ice, nuclei were pelleted at 1,300 g for 4 mins at 4°C, washed with ½ volume PBS, pelleted again, and resuspended in Wash buffer (20 mM HEPES-KOH pH 7.5, 150 mM NaCl, 0.5 mM spermidine, 1x EDTA-free protease inhibitors) at a final concentration of 1 million cells/mL. 5 μL of BioMag Plus Concanavalin A beads (BP531, Bangs Laboratories) per sample were placed on a magnetic stand, liquid removed, and beads washed twice with 1.5 mL binding buffer (20 mM HEPES-KOH pH 7.5, 10 mM KCl, 1 mM CaCl2, 1 mM MnCl2), then resuspended in 5 μL of binding buffer per sample. 100 μL of the nuclei suspension was mixed with 5 μL of equilibrated beads in 0.2 mL PCR tubes and incubated at room temperature for 10 mins. Tubes were placed on a magnetic stand to clear, and liquid was removed. Nuclei-bead complexes were resuspended in 50 μL antibody buffer (Wash buffer with 2 mM EDTA and 0.1% BSA) containing a 1:50 dilution of primary antibodies (H3K27me3, 9733 Cell Signaling Technology; H3K9me3, 39161 Active Motif) or no antibody (IgG control), and incubated on a rotator at room temperature for 1.5 hours. Samples were again placed on a magnetic stand to clear, liquid removed, and resuspended in 50 μL Wash buffer containing a 1:100 dilution of anti-rabbit secondary antibody (ABIN101961, Antibodies-Online). Samples were incubated on a rotator at room temperature for 30 mins, washed with 200 μL Wash buffer, and the liquid discarded. For Tn5 binding, 50 μL of 300-Wash buffer (20 mM HEPES-KOH pH 7.5, 300 mM NaCl, 0.5 mM spermidine, 1x EDTA-free protease inhibitors) with a 1:20 dilution of pAG-Tn5 (15-1017, EpiCypher) was added. After 1 hr at room temperature, nuclei were washed with 300-Wash buffer (once with 200 μL and once with 50 μL) and resuspended in 50 μL tagmentation buffer (300-Wash buffer with 10 mM MgCl2). Tagmentation was performed at 37°C for 1 hr in a thermocycler with a heated lid at 95°C. After tagmentation, samples were placed on the magnet and liquid discarded. Nuclei were resuspended in 50 μL of 10 mM TAPS pH 8.5, placed again on the magnet, and liquid removed. Nuclei were then resuspended in 5 μL release buffer (0.1% SDS, 10 mM TAPS pH 8.5) and incubated at 58°C for 1 hr in a thermocycler with a lid set to 95°C. 15 μL of 0.67% Triton X-100 was added and mixed. For PCR, 2 μL each of 10 μM universal i5 and barcoded i7 primers^70^, and 25 μL NEBNext HiFi 2x PCR Master Mix (NEB M0541S) were added. Cycling conditions were: 58°C for 5 mins, 72°C for 5 mins, followed by 13 cycles of 98°C for 10 sec, 60°C for 10 sec, and 72°C for 10 sec, with a final extension at 72°C for 1 min. Libraries were purified using 1.2x Ampure XP beads and eluted in 16 μL of 10 mM Tris-HCl, pH 8.5. Quality was assessed on a TapeStation D1000 tape, and libraries were pooled for sequencing on a NovaSeq SP 50PE run.

### Nanopore training sample generation

Canonical lambda - Non-methylated Lambda Phage DNA (Sigma-Aldrich, D3654) was used.

CpG lambda genome - Individual 20μL methylation reactions were set up as follows: 1μg Non-methylated Lambda Phage DNA (Sigma-Aldrich, D3654), 20U CpG Methyltransferase M.SssI (NEB, M0226L), 1x NEBuffer 2 (NEB, B7002S), 0.8mM S-adenosylmethionine (SAM) (NEB, B9003S). Samples were incubated for 24hrs at 37°C with an additional 0.8mM SAM spiked in at 4 and 8hrs. Samples were cleaned up as follows: 0.12 U Thermolabile Proteinase K (NEB, P8111S) was added to each reaction and incubated at 37°C for 2hrs then 55°C for 10mins. 22μL (1x volume) H2O was added followed by 44μL (1x volume) AMPure XP beads (Beckman Coulter, A63882). Elution was performed at 50°C for 10mins.

GpC lambda genome - Individual 50μL methylation reactions were set up as follows: 1μg Non-methylated Lambda Phage DNA (Sigma-Aldrich, D3654), 20U GpC Methyltransferase M.CviPI (NEB, M0227), 1x GC Reaction Buffer (NEB, B0227S), 0.8mM SAM (NEB, B9003S). The sample was incubated for 4hrs at 37°C and cleaned up as for the CpG genome.

CpG and GpC lambda genome – the above methods were combined in sequence to methylate both CpG and GpC contexts.

### Long-read ChIMe-seq

gDNA was extracted either as for bulk short reads or using NEB Monarch HMW DNA Extraction for Cells & Blood (T3050L) and agitation of 1400rpm before being quantified with Qubit dsDNA HS Assay Kit (10616763). Library preparation was performed using SQK-LSK109 (Oxford Nanopore) and samples were sequenced on an Oxford Nanopore Promethion 24 with v9.4 flow cells (Oxford Nanopore FLO-PRO002) and MinKNOW (v22.05.10).

### Gastruloid culturing and dissociation

Mouse gastruloids were cultured and dissociated following the protocol previously described^45^ until after the second PBS0 wash post-dissociation.

### FACS testing gastruloid permeabilisation and targeting

ChIMe-seq was performed, in full, as standard, using 0.5 million day 6 mouse gastruloid cells and H3K27me3 antibody as above. Two samples were prepared, one with the standard antibody: H3K27me3 – 1:100 dilution - H3K27me3 Rabbit mAB (C36B11) (CellSignaling, #9733), one using an Alexa Fluor® 647 Conjugate: H3K27me3 – 1:100 dilution - H3K27me3 Rabbit mAB (C36B11) (CellSignaling, #12158S). After the ChIMe-seq stop buffer wash, samples were resuspended in PBS filtered through a 70µm filter (Corning, 431751), into a tube pre-coated with 0.1% ultrapure BSA (ThermoFischer, AM2616) in PBS0, stained with Hoechst 34580 (5 µg/mL) and FACS analysed using a BD Influx cell sorter using a 640nm excitation laser and a 670/30 band pass filter for acquisition.

### Gastruloid ChIMe-seq processing

0.5 million day 5 mouse gastruloid cells were processed with ChIMe-seq as described above but with the addition of 1:40 RNasin Ribonuclease Inhibitor Plus (Promega, N2611) to all buffers.

### Gastruloid ChIMe-seq single cell sorting

After ChIMe-seq cells were resuspended in PBS and stained with Hoechst 34580 (5 µg/mL). Next cells were filtered through a 70µm filter (Corning, 431751) and G1 sorted by FACS using a BD Influx cell sorter into Eppendorf™ twin.tec™ 96 Well LoBind PCR Plates, Skirted (Fischer Scientific, 15280735) containing 2.5µL RLT buffer plus (Qiagen, 1053393). Sorted plates were sealed, centrifuged (2000g, 2mins, 4°C) and stored at −80°C for later use. Remaining unsorted cells were processed as a matched bulk control as for bulk short-read ChIMe-seq.

### scBS-seq library prep

scBS-seq libraries were prepared according to the published protocol^19^ modified to use standard Illumina adapters and improve sequencing quality and alignment rates. Briefly, 96w plates containing sorted cells in 2.5µl of RLT plus buffer were bisulfite converted with the addition of 1.5µl nuclease free water and 26µl of prepared CT conversion reagent (Zymo EZ-Direct, freshly prepared following the manufacturer’s instructions) and incubated at 98°C for 8min followed by 64°C for 3hrs then held at 4°C overnight. Purification was performed by adding 2µl of Zymo Magbeads and 120µl of M-Binding buffer then following the manufacturer’s instructions but with 100µl wash and desulphonation volumes. The converted fragments were eluted in 33.4µl of 1.5µM TruSeqR2_8H oligo (CAGACGTGTGCTCTTCCGATCTHHHHHHHH), heated to 65°C for 5min then cooled to 4°C on a thermocycler. 4µl of 10x buffer (supplied with the klenow exo-), 1.6µl of 10mM dNTPs and 1µl of 50U/µl Klenow exo-polymerase(MCLAB, KPIM-200 or NEB M0212M) was then added and the reaction incubated on a thermocycler using the following program: 4°C 5min, ramping 0.1°C/s to 20°C for 5min then ramping 0.1°C/s to 37°C for 30min. Samples were next heated to 95°C for 1min to denature into single-strands then cooled on ice. 2.5µl of mastermix was added containing 0.25µl of 10x buffer, 0.1µl of 10mM dNTPs, 0.5µl of 10µM TruSeqR2_8H and 0.25µl of 50u/µl klenow exo-then incubated using the same program. This process was repeated another 3 times such that 5 rounds of first strand synthesis had been completed. Samples were purified using 0.7x volumes of SPRI Select beads then to elute were resuspended in 49µl of second strand mastermix containing 1x buffer, 0.4mM dNTPs and 0.4µM of NexteraR1_8N (TCGTCGGCAGCGTCAGATGTGTATAAGAGACAGNNNNNNNN). Without transferring off the beads, samples were heated to 98°C for 2minutes then cooled to 4°C before adding 1µl of 5U/µl klenow exo- and incubating using the same slow ramp program. Following another round of 0.7x SPRI select purification, samples were eluted in 25µl of PCR mastermix (1x KAPA HiFi hotstart readymix, 1µM TruSeq i7 index primer, 1µM Nextera i5 index primer) and cycled as follows: 98°C 45s, followed by 15 cycles of 98°C 15s, 65°C 15s, 72°C 30s. Finally, 5µl of each library was pooled into a 1.5ml eppendorf tube and purified via 2 rounds of 0.7x SPRI select before eluting in 25µl of EB (Qiagen). Samples were sequenced on an Illumina Novaseq6000, NovaseqX or Nextseq2000.

## Data processing

### ChIMe-seq bulk short-read sequencing pre-processing

FastQ sequencing reads were trimmed with trim_galore^71^ (v0.6.7) with the following settings “--paired - -clip_R1 4 --clip_R2 4 --three_prime_clip_R1 4 --three_prime_clip_R2 4 -q 30”. Trimmed reads were aligned to the hg38 reference genome using Bismark v0.23.1^72^ and arguments “bismark -- non_directional -X 2000 --local”. Aligned reads were deduplicated using Bismark “deduplicate_bismark -p –-bam”. Non-bisulfite-converted reads were filtered using a custom processing pipeline detailed at https://github.com/vivekbhr/chime. Methylation information was extracted using Bismark “bismark_methylation_extractor -p --gzip --bedGraph --CX”.

### ChIP-seq pre-processing

Mapped ChIP-seq files used for comparison with ChIMe-seq were downloaded from the ENCODE portal^23^ (accessions: ENCFF392ZKG, ENCFF816ECC, ENCFF897QAS, ENCFF855ZMQ, ENCFF104THG) and processed via the snakePipes ChIP-seq workflow (v1.3.1), with the following parameters: ChIP-seq --singleEnd --fromBAM EXTERNAL_BAM --bamExt .bam –bigWigType log2ratio --bwBinSize 100 -d. -j 20 hg38_mod chip_sample_config.yaml -- local

### WGBS pre-processing

For WGBS-seq, paired-end fastq files obtained from the ENCODE portal^23^ (accession: ENCLB542OXH) were remapped with the same parameters as ChIMe-seq.

### Peak calling

For peak calling, histoneHMM^73^ was used with parameters ‘-b 750 -P 0.1’ for H3K27me3, H3K9me3 and H3K36me3 samples, and MACS2^74^ with parameters ‘-f BAM -g 3091013833 --qvalue 0.001 --keep-dup all --mfold 0 50 –extsize 200’, for H3K4me3 samples. Sample ‘ENCFF226FKB’ was used as control for FDR correction in MACS2.

### Comparison of ChIMe-seq data with other methods

For ChIMe-seq, extracted methylation data was binned into 1kb bins for IGV visualization or 50kb bins for plotting comparison with other methods.

For ChIP-seq, the coverage in 10 and 50kb bins was calculated from the deduplicated BAMs using deepTools^75^ multiBAMSummary with parameters ‘-e 300’. Outlier bin counts (>98th quantile) were removed and the rest of the bins were normalized to counts per million using ‘edgeR::cpm’ function. Bismark’s ‘NoMe_GpC.cov’ output from ‘map_bsChIMÈ workflow was then used to aggregate total and GpC methylation counts per bin onto the same bins, using ‘methylKit::regionCounts’, filtering out bins with less than 5 covered bases^76^. The Pearson correlation was then calculated between the two sets of samples and also between ChIMe replicates on these bins. For WGBS CpG methylation counts were aggregated onto the same bins as above, using ‘methylKit::regionCounts’, filtering out bins with less than 5 covered bases, before calculating Pearson correlation. All samples used for the analysis had a comparable number of reads (80-85 mil), avoiding read-depth related differences.

### Specificity and sensitivity analysis

For specificity analysis of ChIP-seq samples deduplicated bam files were intersected with the relevant peak file using bedtools intersect. The number of reads in the intersected bam / total number of reads in the deduplicated bam provided the fraction of reads in peaks. For ChIMe-seq samples the 3-base context of a cytosine was identified from the reference and the result intersected with the relevant peak file as above. Total GCH calls and GCH methylation calls genome wide and in peaks were counted. In peak GCH methylation calls / genome wide GCH methylation calls provided the fraction of methylation in peaks. GCH methylation calls / total GCH calls in peaks provided GpC methylation fraction within peaks.

### CUT&Tag analysis

Raw reads were demultiplexed using i7 barcodes and trimmed to remove transposase sequences using Cutadapt^77^. Reads were aligned to the mm39 reference genome with Bowtie2^78^ using the parameters - -local --quiet --very-sensitive-local --no-mixed --no-discordant --phred33 -I 10 -X 700, then sorted and indexed using samtools^79^. Duplicate reads were removed using Picard MarkDuplicates^80^ with -- REMOVE_DUPLICATES true.

Aligned BAM files were converted to bigWig format for IGV visualization using bamCoverage^75^ with parameters -bs 1 --normalizeUsing RPKM. Read counts for 50 kb genomic windows were computed using multiBamSummary^75^ in BED-file mode with parameters -e 300 --outRawCounts -- smartLabels. The BED file contained 50 kb genomic bins generated by methylKit::regionCounts^76^ for ChIMe-seq samples. For downstream correlation analysis, IgG signal was averaged across replicates, and the top 15% of bins with the highest IgG signal were excluded to remove open chromatin background characteristic of Tn5-based methods. Outlier bins (>98th percentile) were also removed. The remaining bins were normalized to counts per million (CPM) using edgeR::cpm^81^. Pearson correlations were then calculated between CUT&Tag and ChIMe-seq samples, as well as between ChIMe-seq replicates.

### Retraining of ONT models

Three samples were used for training the CpG and GpC models. One canonical, unmethylated Lambda phage sample and two enzymatically methylated to near completion in the CpG or GpC context.

For each sample, Megalodon (https://github.com/nanoporetech/megalodon; v2.5.0) was run on a subset of the data for its signal_mappings output, using Guppy (v5.0.11), marking reference motifs CG or GC as modified where relevant. As modifications may influence basecalling qualities, reads with many modifications may fall into the fail set. Therefore, reads in the fail dataset were included for training, in proportion to their occurrence in the full dataset. The signal_mappings output data was merged into a single file using Taiyaki merge (https://github.com/nanoporetech/taiyaki; v5.3.0). The merged data was prepared for training using Remora (https://github.com/nanoporetech/remora) dataset prepare, specifying the motif as ‘HCG’ (for CpG model) (v2.1.0) or ‘GCH’ (for GpC model) (v1.1.1) with the modification at the C position for each.

Models were trained using Remora model train for each motif. For the CpG model an existing Remora model was finetuned. As the version of Remora used (v1.1.1) did not support finetuning existing models, nor include checkpoint models, the CpG model from Remora v2.1.0 (dna_r9.4.1_e8_hac_v3.5.1_5mc_CG_v0.chk), which did not support the fast5 data used, was backported. This model was loaded as a checkpoint and trained further with a reduced learning rate (‘- -lr 0.0001’). The GpC model was trained from scratch.

For training both models the number of ‘epochs’ was set to 250 with ‘early-stopping’ at 10, causing training to end when no measurable improvements were made for 10 epochs in a row. The CpG model ended training after 59 epochs, while the GpC model ended training after 24 epochs. The ‘model_best’ was selected for each, as the ‘final_model’ showed no improvements but could have introduced some overtraining.

### Long-read mapping and modification calling

For GpC methylation modification calling, the fast5 reads were processed via Megalodon (https://github.com/nanoporetech/megalodon; v2.5.0) using guppy basecaller backend (v4.2.2, using cuda 11) with the following parameters: ‘--guppy-config dna_r9.4.1_450bps_hac.cfg --guppy-timeout 500 --reference hg38.fa --outputs mod_mappings mods per_read_mods --mod-output-formats bedmethyl modvcf wiggle --write-mods-text --write-mod-log-probs --remora-model retrained_model.onnx’ where ‘retrained_model.onnx’ file was obtained from the above retraining. Next the methylation status of each site in per_read_modified_base_calls.txt files was binarised at a mC/C probability threshold of >0.8 and otherwise discarded. For CpG calls the 4-base NNCG context was obtained from the reference and CACG, TCCG, ACCG and CCCG contexts were removed.

### Comparison to matched Illumina sample

Following ChIMe-seq targeting of H3K27me3 both bulk short-read ChIMe-seq and long-read ChIMe-seq library preparations were performed on the same cells. Both CpG and GpC data types were processed as above with 1kb bins used for IGV plots and 50kb bins used for scatter plots. Surrounding NNCGN contexts were extracted from reference for CG calls made from Illumina data and Nanopore data called with Remora v1.1.1, v2.1.0 and our finetuned model. Average methylation rates for each context were compared with matched Illumina data as ground truth. Comparison of calls within and outside H3K27me3 domains, called from sortChIC data^18^ as above for ChIP-seq data, was made. Domain overlap of CpG calls was determined using bedtools intersect. Absolute (ONT − Illumina) and relative ((ONT − Illumina)/Illumina)) differences in Nanopore data were assessed per context.

### Single-read CpG Vs GpC comparison

From processed per_read_modified_base_calls.txt files per-read GCH and CGH methylation means were calculated and the individual summaries were joined before plotting.

### Single-molecule CpG and GpC modification visualization

Genome browser aggregate tracks were generated using 1kb binned bigwigs, as above for IGV, and the WashU Epigenome Browser. Single-molecule browser tracks for both modifications were generated as in DiMeLo-seq^13^.

### Comparison of Nanopore and Illumina methylation results by context

For each caller files were filtered to contain only chromosome 2 and methylation calls were binarised at mC or C probability 0.8 from per_read_modified_base_calls.txt files for Nanopore samples and using Bismark coverage outputs for Illumina control. NHCGN contexts were added from reference. Methylation averages were calculated per context and across chromosome and plotted. Methylation calls were split into inside/outside GpC methylation domains using bedtools intersect and peaks called on sortChIC data as above. Absolute differences for each NHCGN context were (ONT average methylation) – (Illumina average methylation) per context and region. Relative difference was calculated as (ONT average methylation) / (Illumina average methylation). Samples had contexts filtered and this analysis was repeated. Bulk QCs of methylated lambda training sets were pre-processed as for bulk short-read ChIMe-seq samples then treated as an Illumina sample here.

### In silico fragmentation of Nanopore reads

Start locations of reads mapped with CpG and GpC workflows were extracted using samtools^79^ view ${1} | awk -F ‘\t’ ‘{print $1, $3, $4, $9}’ > cut_read_start_${1}.txt. All reads were then checked for mapping to the same location in both workflows and discarded if not. Reads were fragmented to 1000bp by appending a numeric suffix to Read IDs based on each modification calls relative position from the start of the read as a multiple of 1000. Summaries of CpG methylation statistics were generated from the data by dividing methylated CpGs per fragment by total CpGs per fragment. Individual fragments were filtered to retain only those exactly 1000bp long, and containing at least 3 CpGs. Fragments were partitioned into separate files based on their CpG methylation fraction.

### Shuffling of methylation status

Individual methylation statuses were shuffled by randomly sampling methylation statuses without replacement across the sample. This output was then fed to fragmentation and downstream analysis as above.

### Computing GpC methylation conditional probability curves from processed Nanopore reads

Pairwise GpC methylation relationships were calculated for fragments within each CpG methylation interval and histone-modification dataset. Within individual 1-kb fragments, each methylated GpC was used as an anchor, and the absolute genomic distance to every other called GpC within the same fragment was determined. GpC pairs were classified as methylated–methylated (MM) when both the anchor and comparison GpC were methylated, or methylated–unmethylated (MU) when the anchor GpC was methylated and the comparison GpC was unmethylated. Pair counts were pooled across fragments within each group. For each genomic separation, *d*, the conditional probability that a GpC was methylated given that another GpC (d) base pairs away was methylated was calculated as

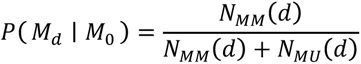

where *N*_MM_ (*d*) and *N*_MU_ (*d*) denote the numbers of methylated–methylated and methylated–unmethylated GpC pairs, respectively, separated by *d* bp. Distances were evaluated at single-base-pair resolution without additional distance binning.

### Mathematical model of nucleosome organization

Chromatin was modelled as a one-dimensional sequence of alternating nucleosome-protected and linker regions. To derive the spatial correlations generated by this model, we adapted a transfer-matrix approach previously developed for nucleosome positioning^82^. Unlike the original formulation, in which nucleosome spacing is determined by a specified interaction energy, the transfer matrix was constructed directly from arbitrary probability distributions of nucleosome footprint and linker lengths. This formulation therefore does not assume a particular physical interaction between neighbouring nucleosomes. For the analyses presented here, nucleosome footprint and linker lengths were modelled using gamma distributions.

The transfer matrix describes transitions between nucleotide-level sub-states representing positions within nucleosome-protected and linker regions. Transition probabilities were determined directly from the specified nucleosome footprint and linker length distributions. The analytical transfer-matrix solution was then used to calculate the pairwise conditional probability of nucleosome occupancy as a function of genomic distance.

Because ChIMe-seq measures GpC methylation rather than nucleosome occupancy directly, an observation model was incorporated that accounts for different GpC methylation probabilities within nucleosome-protected and linker regions. The finite lengths of histone modification domains was additionally accounted for. In the observation model, these domains represent genomic regions in which methylated GpCs can be detected, whereas outside these regions they cannot. As a result, a methylated anchor GpC must lie within a modification domain. At increasing distances from the anchor, a second GpC may either remain within the same domain, in which case its methylation probability is determined by the nucleosome model, or lie outside the domain, in which case it is assumed to be unmethylated. Exit from a domain was modelled as a memoryless process with a constant probability (λ) per unit bp, resulting in an exponential decay of the conditional methylation signal with distance.

The resulting model relates nucleosome footprint and linker length distributions, their respective GpC methylation probabilities, and the domain boundary density parameter (λ) to the conditional GpC methylation profiles measured experimentally. Model parameters were inferred by minimizing the least-squares difference between the predicted and measured conditional GpC methylation profiles. Full mathematical derivations of the transfer matrix, observation model, and modification-domain are provided in Supplementary Note.

### Model fitting and optimization

Model parameters were estimated by minimizing the residual sum of squares (RSS) between the predicted and observed conditional GpC methylation curves. Optimization was performed using the differential-evolution algorithm implemented in SciPy (scipy.optimize.differential_evolution). Differential evolution is a population-based stochastic global optimization algorithm and was used to reduce sensitivity to parameter initialization and local minima. For each oscillation curve, the optimization was performed independently 10 times, and the parameter set from the run that exhibited the lowest RSS was kept as the final solution.

The mean nucleosome-protected length was fixed at 147 bp. Six parameters were estimated: the mean linker length, the standard deviations of the nucleosome-protected and linker-length distributions, the domain boundary density (λ), and the methylation probabilities (P_linker_) and (P_nuc_). The mean linker length was constrained to 20–80 bp, both length-distribution standard deviations to 10–60 bp, λ to 0–0.001, P_linker_ to 0–1, and P_nuc_ to 0–0.5.

For numerical evaluation of the nucleosome model, the gamma-distributed state lengths were truncated to obtain a finite state space. For each distribution, the state space was extended to the first integer genomic distance at which its cumulative probability exceeded 0.99. The resulting finite transition matrix describes progression through nucleosome-protected and linker states. Distance-dependent conditional state probabilities were calculated by iteratively propagating the probabilities of individual nucleosome and linker substates, accounting for the probability of remaining within a region or transitioning to the next. This procedure is mathematically equivalent to multiplication by the transition matrix but avoids explicitly constructing and diagonalizing it. These calculations and steps were repeated for each candidate parameter set during differential-evolution optimization. Further details on model fitting and optimization are provided in Supplementary Note.

### Bulk gastruloid control data processing

Bulk gastruloid data was processed as for bulk short-read sequencing except data was mapped to the mouse mm10 reference genome.

### scBS-seq data preprocessing

Read 1 only of paired-end sequencing reads for each single cell were trimmed with trim_galore to remove low quality read ends and N sequences using arguments --2colour 30 and --trim-n. Reads were then hardclipped by 9bp at 3’ end and 19bp at 5’ end before removing reads with average phred score < 30 and final length <20bp using cutadapt. Reads were mapped to the mouse mm10 reference genome with Bismark using ‘bismark --non_directional --se’. Aligned reads were deduplicated using Bismark and default settings for single-end reads. Deduplicated reads were filtered for bisulfite underconversion using Bismark filter_non_conversion with a required minimum of 5 non-CpG cytosines and a cutoff of 50% non-CpGme. Methylation information was extracted from filtered bams with Bismark in --CX mode ‘bismark_methylation_extractor -s --gzip --bedGraph --CX’. Cytosine reports were generated with ‘coverage2cytosine --nome-seq’ in order to be GpCme aware. Strand information was removed by converting all minus strand methylation calls to their respective plus strand positions and methylation calls were binarised per-position as described previously^57^.

### scBS-seq gastruloid pseudobulk signal enrichment analysis

H3K27me3 domains were detected for day 5 gastruloid T-ChIC data using histoneHMM with parameters ‘-b 750 -P 0.1’. Enrichment plots were made using deeptools (v3.5.1) computeMatrix scale-regions and computeMatrix reference-point.

### scBS-seq global metrics

Per-cell methylation, motif observation and read counts were extracted from Bismark coverage files and summary reports using custom scripts detailed in https://github.com/ejolysmith/ChIMe-seq.

### Single-cell CpG and GpC methylation analysis and cell clustering

CpG and GpC methylation calls were analysed independently using MethSCAn^83^, following the standard MethSCAn workflow with an addition of filtering for cells with > 100,000 CpGs covered and < 3% HCHme. For each modality, methylation profiles were smoothed using methscan smooth. Variably methylated regions (VMRs) were identified using methscan scan with a bandwidth of 1000 bp. Cell-by-VMR matrices were generated using methscan matrix, and mean shrunken residuals were used for downstream analysis. VMRs measured in fewer than 15% of cells were excluded. Missing values were imputed using a Python implementation of the iterative PCA procedure described in the MethSCAn workflow. Features were mean-centered and missing values were initially set to zero. Missing values were then iteratively replaced with values reconstructed from the principal components until convergence.

For joint analysis, the filtered CpG and GpC residual matrices were concatenated before iterative PCA. Twelve principal components were retained. Leiden clustering was performed in this space using a 10-nearest-neighbour graph based on correlation distance and a resolution of 0.43. Cells were visualized using UMAP with 20 nearest neighbours, a minimum distance of 0.2, and correlation distance. For comparison, CpG and GpC matrices were also analysed separately. Iterative PCA was performed for each modality, and 20 principal components were retained. CpG-only and GpC-only UMAPs were generated using 20 nearest neighbours, a minimum distance of 0.2, and correlation distance.

To identify cluster-associated methylation differences, each joint Leiden cluster was compared with all other cells in a one-versus-rest analysis. methscan diff was run separately on the CpG and GpC datasets using a bandwidth of 1000 bp.

### T-ChIC data pre-processing, cell typing and lineage assignment

T-ChIC data was processed as previous^45^.

### T-ChIC H3K27me3 differential enrichment analysis

T-ChIC H3K27me3 counts^45^ were quantified over genomic regions corresponding to the ChIMe-seq GpC differentially methylated regions (DMRs). For each lineage, cells were grouped by biological replicate and H3K27me3 counts within each DMR were summed to generate pseudo-bulk samples. The lineage of interest was compared with a pooled reference comprising all remaining lineages from the same replicate. Differential H3K27me3 enrichment was calculated using PyDESeq2 with lineage grouping as the design factor and the pooled remaining lineages as the reference. Pluripotent, ectoderm, mesoderm and endoderm cells were each analysed independently in a one-versus-rest comparison.

### T-ChIC RNA-seq differential expression analysis

Lineage-specific differential gene expression was calculated from the RNA component of the T-ChIC gastruloid dataset^45^. Day-5 gastruloid cells were grouped by lineage, and each lineage was compared with all remaining cells in a one-versus-rest analysis. For each gene, log2 fold change was calculated as the difference between the log2-transformed mean expression in the target lineage and the remaining cells, using a pseudocount of 0.1. Statistical significance was assessed using Welch’s two-sample t-test, and P values were corrected for multiple testing using the Benjamini–Hochberg procedure. Log2 fold changes and adjusted P values were used for comparison with ChIMe-seq methylation profiles.

### Comparison of ChIMe-seq single-cell clusters with T-ChIC chromatin and transcriptomic profiles

To compare ChIMe-seq GpC methylation with T-ChIC H3K27me3 enrichment, measurements from the two assays were matched using the genomic coordinates of the GpC DMRs. ChIMe-seq DMRs were retained at an absolute methylation difference ≥0.05 and unadjusted P < 0.05, and T-ChIC regions were retained at an absolute log2 fold change ≥0.5 and unadjusted P < 0.05. Correlations between ChIMe-seq GpC methylation differences and T-ChIC H3K27me3 log2 fold changes were calculated for each cluster–lineage pair.

To compare ChIMe-seq signals with lineage-specific gene expression, CpG and GpC DMRs were assigned to the nearest transcription start site (TSS) using mm10 mouse gene annotations from the UCSC Genome Browser (knownGene and kgXref tables). For CpG methylation, DMRs with midpoints located between 1 kb upstream and 500 bp downstream of the nearest TSS were retained, accounting for gene strand. For the H3K27me3-associated GpC signal, DMRs overlapping a 4-kb window on either side of the nearest TSS were retained. ChIMe-seq DMRs were filtered for an absolute methylation difference >0.05 and unadjusted P < 0.05. Lineage-specific differentially expressed genes were filtered for an absolute log2 fold change >1 and an adjusted P < 0.05. Where multiple DMRs were assigned to the same gene, methylation differences were averaged. Correlations between differential methylation and RNA log2 fold change were calculated for each ChIMe-seq cluster and T-ChIC lineage pair.

ChIMe-seq clusters were assigned to lineages based on their correspondence with lineage-specific H3K27me3 and RNA expression profiles. The resulting lineage proportions were compared with those reported for day 5 and day 6 T-ChIC gastruloids. Ninety-five per cent confidence intervals for ChIMe-seq lineage proportions were estimated from 1000 multinomial bootstrap samples.

### Computing GpC methylation conditional probability curves from single-cell ChIMe-seq data

GpC methylation conditional probability curves were calculated independently for each cell using the approach described above. Binarised GpC methylation calls from the scBS-seq pre-processing were used. Pairs from the upstream and downstream directions were pooled, and conditional probabilities were calculated at distances from 3 to 999 bp. Only called GpC sites contributed to the pair counts. No smoothing was applied to the resulting single-cell profiles.

### Fitting single-cell GpC methylation conditional probability curves

For single-cell data, the nucleosome-linker model was extended to account for GpC observations originating from different homologous chromosomes. Two GpCs contributing to a conditional methylation measurement can originate from the same or different homologues, each with probability 0.5. We introduced an inter-allelic concordance parameter, *k*, defined as the probability that a genomic position is in a linker state on one homologue given that the corresponding position is in a linker state on the other. The contribution of different-allele pairs to the nucleosome-dependent oscillations was incorporated through an amplitude-scaling parameter, 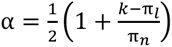, where π_l_ = µ_l_/ (µ_n_ + µ_l_) and π_n_ = µ_n_/ (µ_n_ + µ_l_) are the mean linker and nucleosome occupancies, respectively. When nucleosome–linker organization is independent between homologues, *α* = 0.5 whereas perfect concordance gives *α* = 1. Single-cell curves were initially fitted with all seven model parameters allowed to vary, with *α* constrained between 0.5 and 1. The mean fitted value of *α* across cells was 0.795, corresponding to an inter-allelic concordance probability of approximately *k* = 0.7 for the inferred linker lengths. Cells were subsequently refitted with *α* fixed at 0.795 to reduce the number of free parameters and improve estimation of the remaining parameters. The mathematical derivation is provided in Supplementary Note.

### Analysis of nucleosome organization by CpG methylation state from single-cell ChIMe-seq data

CpG methylation profiles were aggregated separately for each lineage. The genome was divided into non-overlapping 1-kb bins. For each bin, the CpG methylation fraction was calculated from all methylated and unmethylated CpG calls across cells. Bins containing fewer than three detected CpG positions were excluded. The remaining bins were divided into ten CpG methylation groups from 0 to 1 in increments of 0.1. To estimate uncertainty, cells within each lineage were resampled with replacement 100 times. Each bootstrap sample contained the same number of cells as the original lineage. CpG methylation fractions were recalculated for each bootstrap sample, and genomic bins were reassigned to CpG methylation groups.

GpC conditional methylation probability curves were then calculated separately for each CpG methylation group. Only GpC observations within genomic bins assigned to that group were included. GpC pairs were formed only between observations from the same cell. Pair counts were pooled across cells according to their bootstrap sampling weights. Conditional probability curves were fitted using the nucleosome-linker model described above, with the chromosome-synchronization parameter *α* fixed to 0.795. Model parameters were estimated independently for each lineage, CpG methylation group and bootstrap sample. Mean parameter estimates and standard deviations were calculated across the 100 bootstrap samples.

## Data availability

Raw and processed data of this study are submitted to Gene Expression Omnibus (GEO) and available under accession numbers GSE345419 (CUT&Tag), GSE345431 (4T1 cells), GSE345435 (bulk ChIMe-seq), GSE345655 (nanopore data), GSE345447 (single-cell data). Public data used in this study can be found under K562 bulk ChIP data and WGBS (H3K4me3, ENCFF855ZMQ; H3K9me3, ENCFF104THG; H3K27me3, ENCFF392ZKG; H3K36me3, ENCFF816ECC; ChIP input control, ENCFF897QAS; WGBS, ENCLB542OXH). T-ChIC data is available as reported^45^.

## Code availability

Pre-processing scripts for bulk short-read and long-read data can be found at https://github.com/vivekbhr/chime. All other scripts can be found at https://github.com/ejolysmith/ChIMe-seq

## Acknowledgements

The authors thank the Hubrecht Sorting Facility (HSP), Utrecht Sequencing Facility (USEQ) and Anders Didriksen of the Protein Facility at the Hubrecht Institute for cell sorting, sequencing and assistance with protein purification respectively. J.V-L. was supported by Horizon 2020 Marie Skłodowska-Curie Actions Initial Training Network “EpiSyStem” (grant number 765966). E.J. was supported by the NWO Summit Grant “Evolving life from non-life (EVOLF)” (SUMMIT.1.004). The SNF (P2BSP3-174991), HFSP (LT000209/2018-L) and Marie Skłodowska-Curie Actions (798573) supported P.Z. The EMBO LTF (ALTF 1197–2019) supported V.B. S.T-G. was supported by The Branco Weiss Fellowship–Society in Science, administered by ETH Zürich. This work was supported by a European Research Council Advanced grant (ERC-AdG grant no. 101053581-scTranslatomics). This work is part of the Oncode Institute, which is partly financed by the Dutch Cancer Society. The funders had no role in study design, data collection and analysis, decision to publish or preparation of the manuscript. Schematic illustrations were created in BioRender.

## Contributions

J.V-L., P.Z. and A.v.O. designed the project; J.V-L. and P.Z. developed laboratory techniques; E.J-S. developed and applied the chromatin model; J.V-L. performed experiments; J.V-L and E.J-S. analysed the data; F.M. assisted with enzyme purification, V.B. processed ENCODE data and pre-processed bulk ChIMe-seq data; R.S. developed Oxford Nanopore models; S.M. performed exploratory non-model NRL calculations; M.B. cultured gastruloids; P.Z. and M.B. analysed T-ChIC data; S.C. performed scBS-seq; S.T-G. performed external experiments. W.R. and J.d.R. provided supervision. J.V-L. E.J-S. and A.v.O. wrote the manuscript.

## Author information

### Current Addresses

**Princess Máxima Center for Pediatric Oncology, Utrecht, the Netherlands.**

Joe Verity-Legg, Roy Straver, Jeroen de Ridder

**Department of Molecular Biology and Genetics, Aarhus University, Aarhus, Denmark.**

Marloes Blotenburg, Peter Zeller

**Institute of Biodynamics and Biocomplexity, Dept. of Biology, Utrecht University, the Netherlands.**

Vivek Bhardwaj

## Ethics declarations

ChIMe-seq is the subject of a patent application GB2626748.4 on which J.V-L., S.T-G. and A.v.O. are inventors. J.d.R. is cofounder and shareholder of Cyclomics B.V.. S.J.C., and W.R. are employees of Altos Labs. All other authors declare no competing interests.

---

**Supplementary Figure 1.**
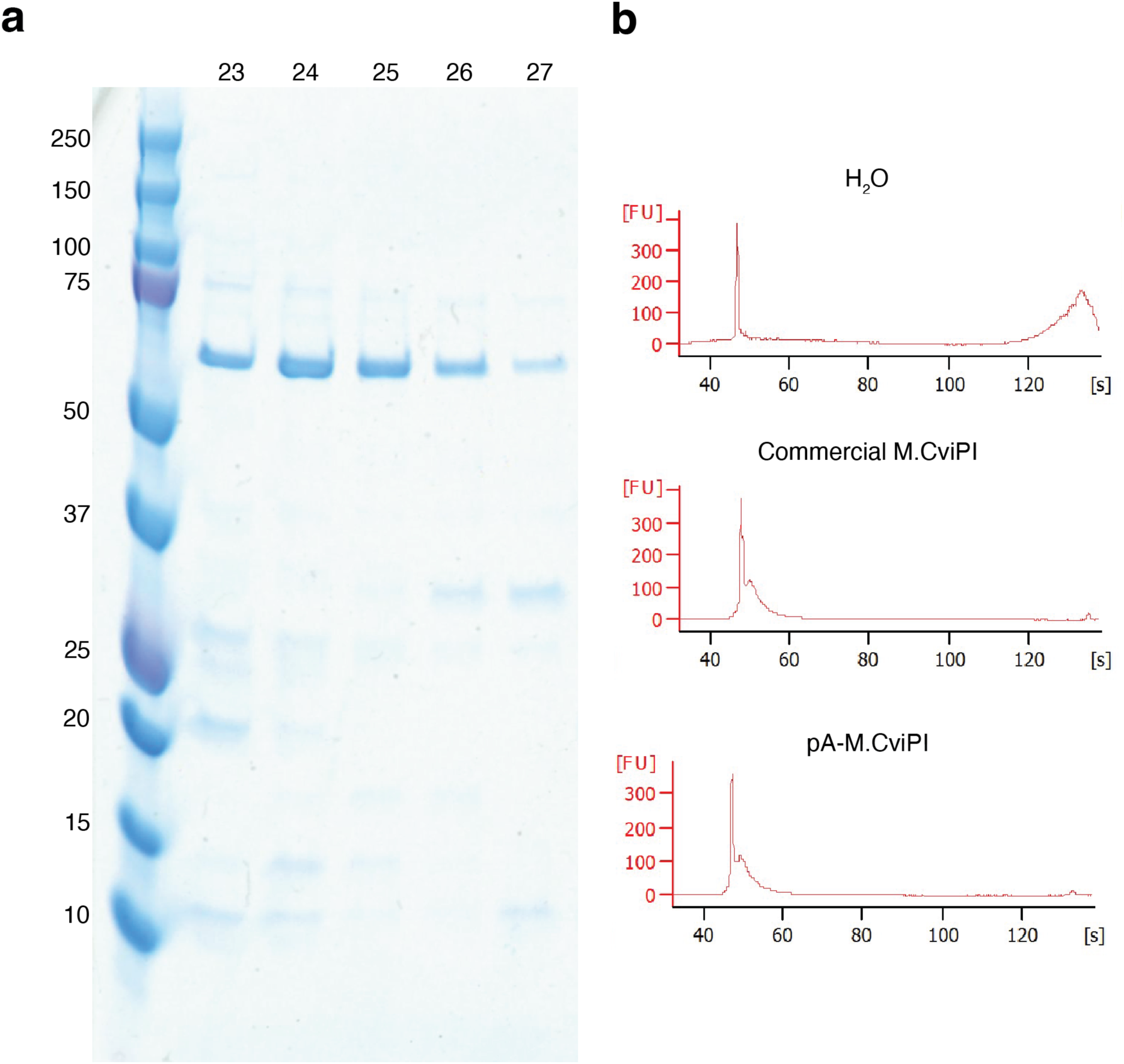
Expression, purification, and enzymatic validation of pA-M.CviPI. **a.** Coomassie-stained polyacrylamide gel demonstrating the successful purification of the 63-kDa pA-M.CviPI fusion protein. Gel lane numbering indicates the sequentially collected high-performance liquid chromatography (HPLC) eluate fraction, y-axis numbering indicates kDa values of the protein ladder. **b.** Bioanalyzer traces showing the effect of methylation-dependent MspJI digestion on unmethylated lambda phage genomic DNA following treatment with H₂O (negative control), commercial M.CviPI (positive control), or pA-M.CviPI.

**Supplementary Figure 2.**
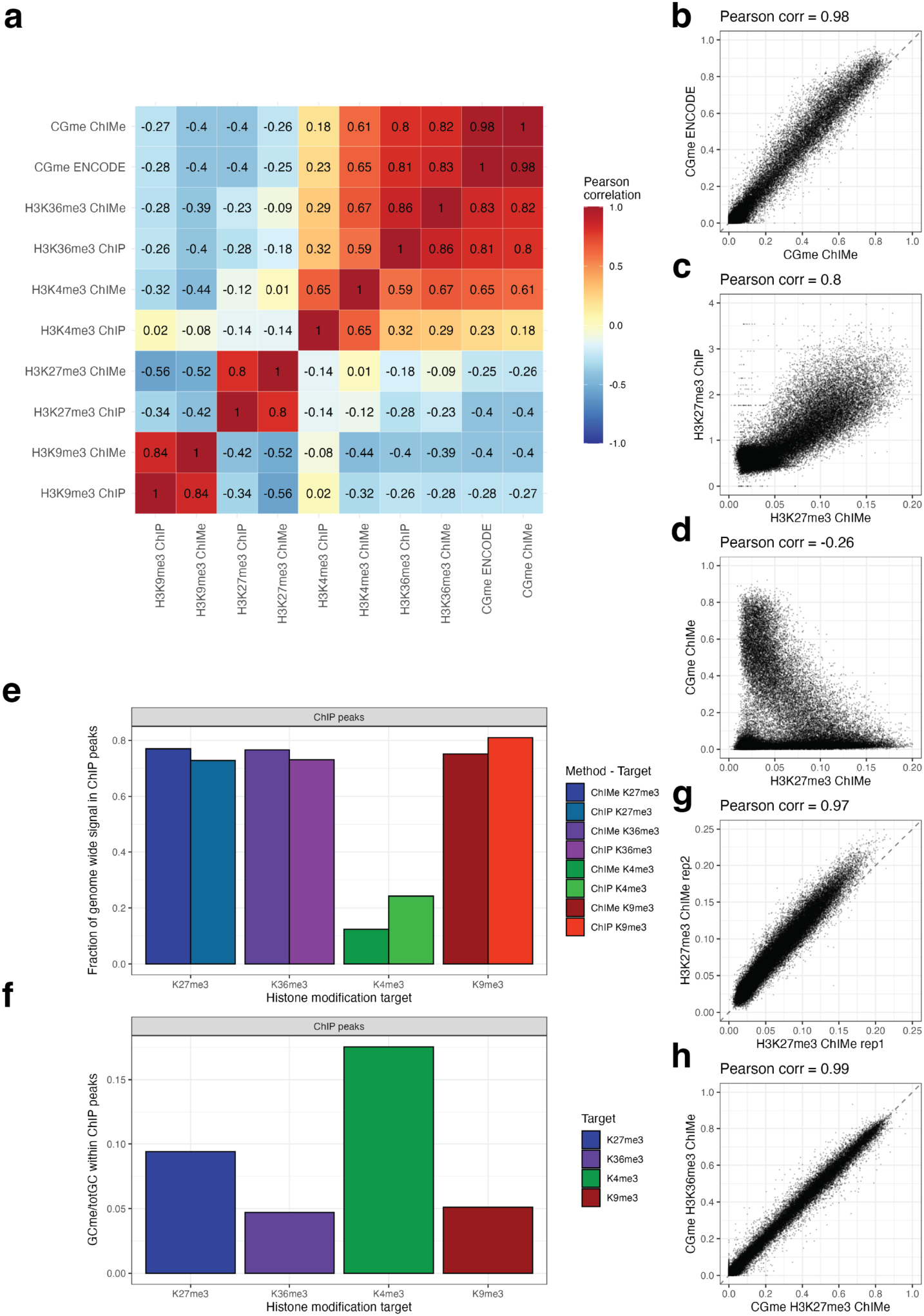
Quantitative validation of ChIMe-seq accuracy, specificity, and reproducibility. **a.** Heatmap showing Pearson correlation coefficients between all samples displayed in Fig. 1b. Correlations were calculated using 50-kb genomic bins. **b.** CpG methylation levels measured by ChIMe-seq versus whole-genome bisulfite sequencing (WGBS). **c.** ChIMe-seq GpC methylation levels versus ChIP-seq normalized read counts for H3K27me3. **d.** CpG methylation levels versus GpC methylation levels from a single H3K27me3-targeted ChIMe-seq experiment. **e.** Fraction of all genome-wide, immuno-targeted signal (ChIMe-seq; individual GpCme calls, ChIP-seq; sequencing read counts) overlapping peaks called from ENCODE ChIP-seq data. **f.** Fraction of individual GpC motif observations within ENCODE ChIP-seq peaks called as methylated. **g.** Comparison of GpC methylation levels between biological replicates targeting the same histone modification. **h.** Comparison of CpG methylation levels between ChIMe-seq experiments targeting different histone modifications.

**Supplementary Figure 3.**
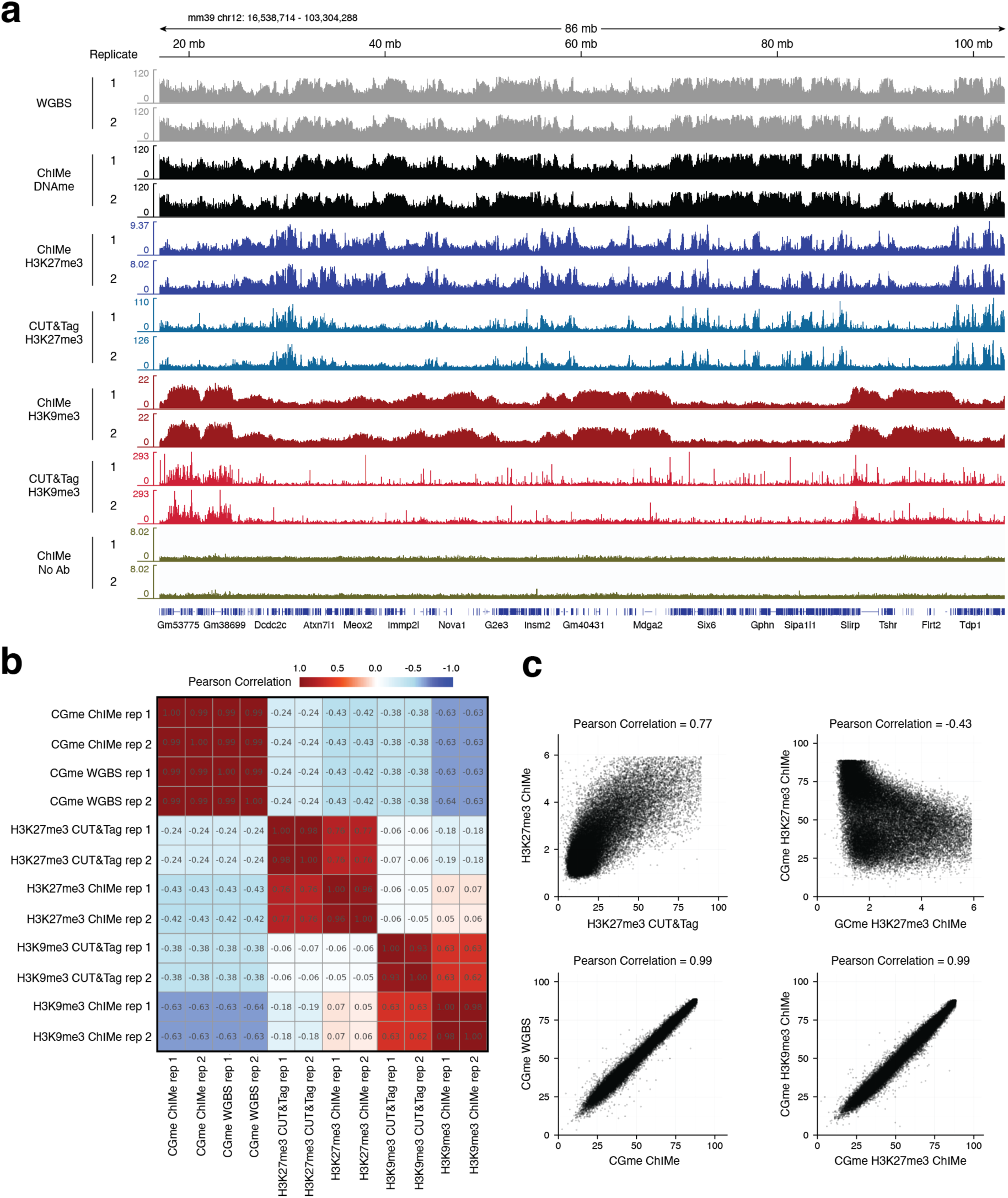
Independent validation of ChIMe-seq performance in a second laboratory. **a.** Genome browser tracks showing ChIMe-seq profiles for multiple histone modification targets alongside corresponding CUT&Tag and whole-genome bisulfite sequencing (WGBS) datasets generated from the same 4T1 cell culture. The ChIMe-seq DNA methylation (DNAme) tracks are derived from the H3K27me3-targeted experiments. All tracks are displayed in 1-bp bins, visualised with mean windowing in IGV. y-axes indicate DNA methylation levels for methylation-based measurements and read counts for CUT&Tag datasets. **b.** Heatmap showing Pearson correlation coefficients between all samples displayed in **a**. Correlations were calculated using 50-kb genomic bins. **c.** Representative scatter plots underlying the correlation analyses shown in **b**, calculated in 50-kb bins.

**Supplementary Figure 4.**
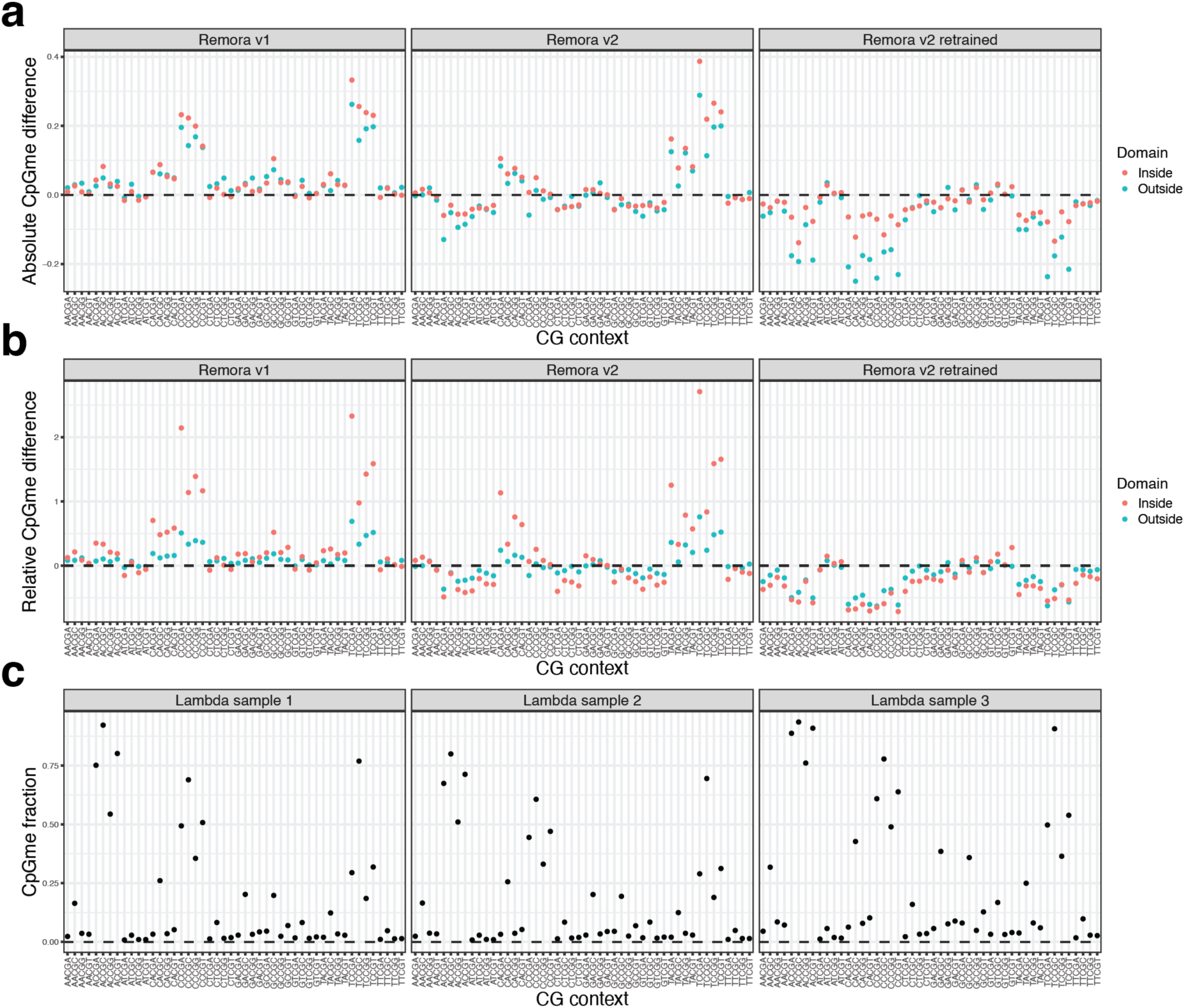
Sequence-context optimization of ONT CpG methylation calling. **a.** Difference in average CpG methylation levels between ONT and Illumina sequencing for each NHCGN 5-mer sequence context on chromosome 2, for a single H3K27me3-targeted ChIMe-seq sample. Difference is calculated as ONT CpGme − Illumina CpGme. Dashed line indicates parity with Illumina results. Non-zero values represent a deviation from the expected result based on Illumina sequencing. Positive values indicate higher methylation levels in ONT results. Facets correspond to the three ONT methylation callers, including our retrained model. Point colour distinguishes sites located within and outside H3K27me3/GpC methylation domains called from sortChIC data^18^. **b.** As in **a**. but for relative difference in CpG methylation levels between ONT and Illumina sequencing, calculated as (ONT CpGme – Illumina CpGme) / Illumina CpGme. **c.** CpG methylation levels for each 5-mer sequence context measured from Illumina sequencing of lambda phage DNA treated with M.CviPI to generate a GpC only methylation sample as training data. Reproducible CpG methylation is observed across three biological replicates, after treatment with GpC methyltransferase.

**Supplementary Figure 5.**
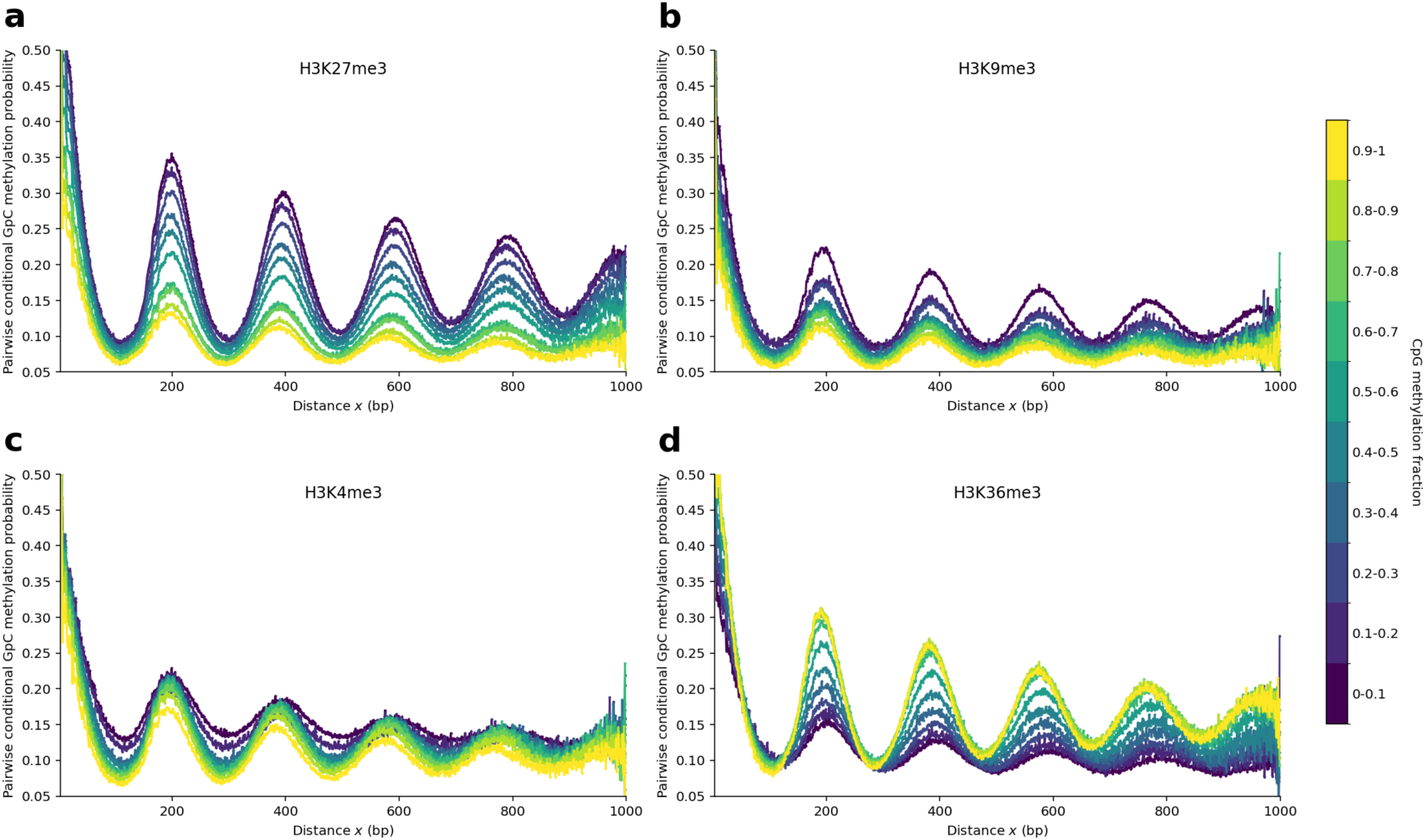
Pairwise conditional GpC methylation oscillations across histone modifications and CpG methylation fractions. **a–d.** Measured pairwise conditional GpC methylation probability as a function of genomic distance x for H3K27me3 (**a**), H3K9me3 (**b**), H3K4me3 (**c**), and H3K36me3 (**d**). Each curve corresponds to a fragment group binned by CpG methylation fraction, with colours indicating the methylation bins.

**Supplementary Figure 6.**
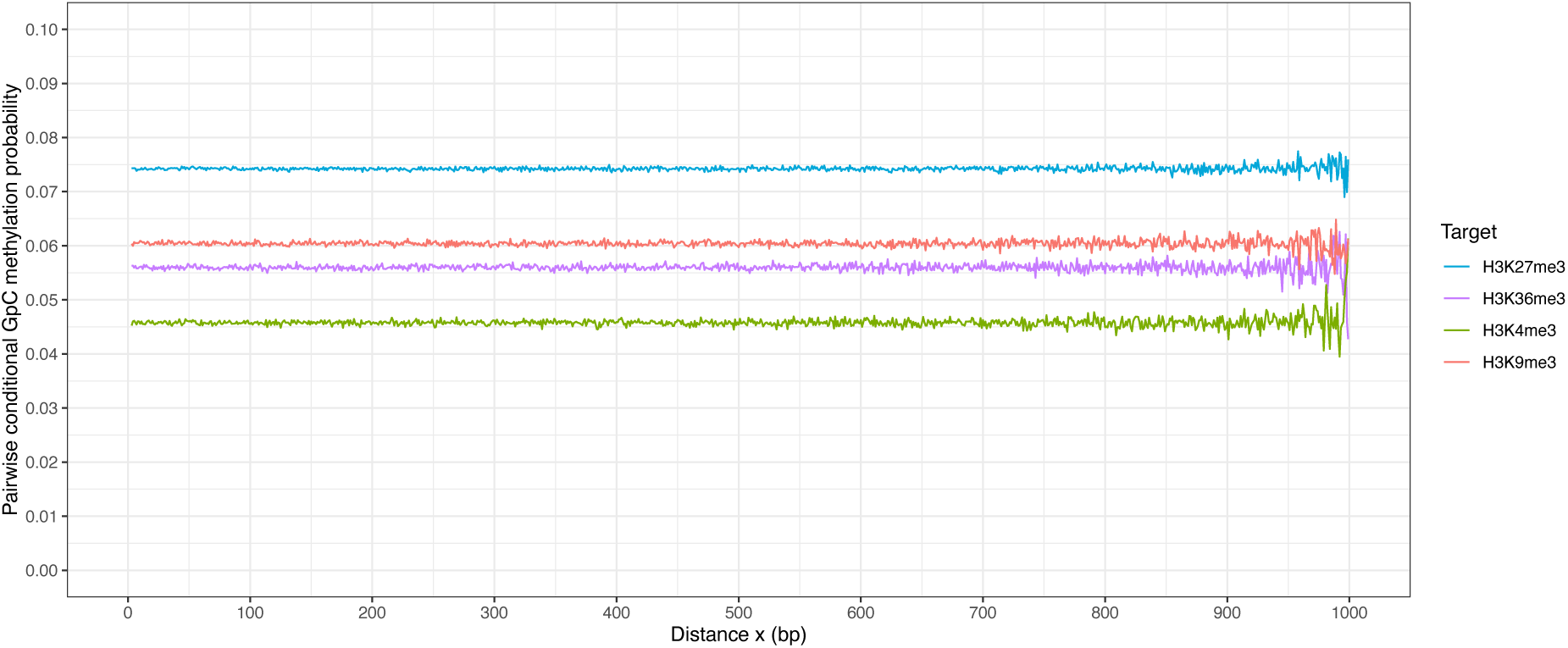
Randomization of GpC methylation removes conditional methylation oscillations. Pairwise conditional GpC methylation probability as a function of genomic distance for ChIMe-seq experiments targeting different histone modifications, after randomly shuffling GpC methylation states across the sample while preserving GpC positions. Curves show the fragment group containing no methylated CpGs.

**Supplementary Figure 7.**
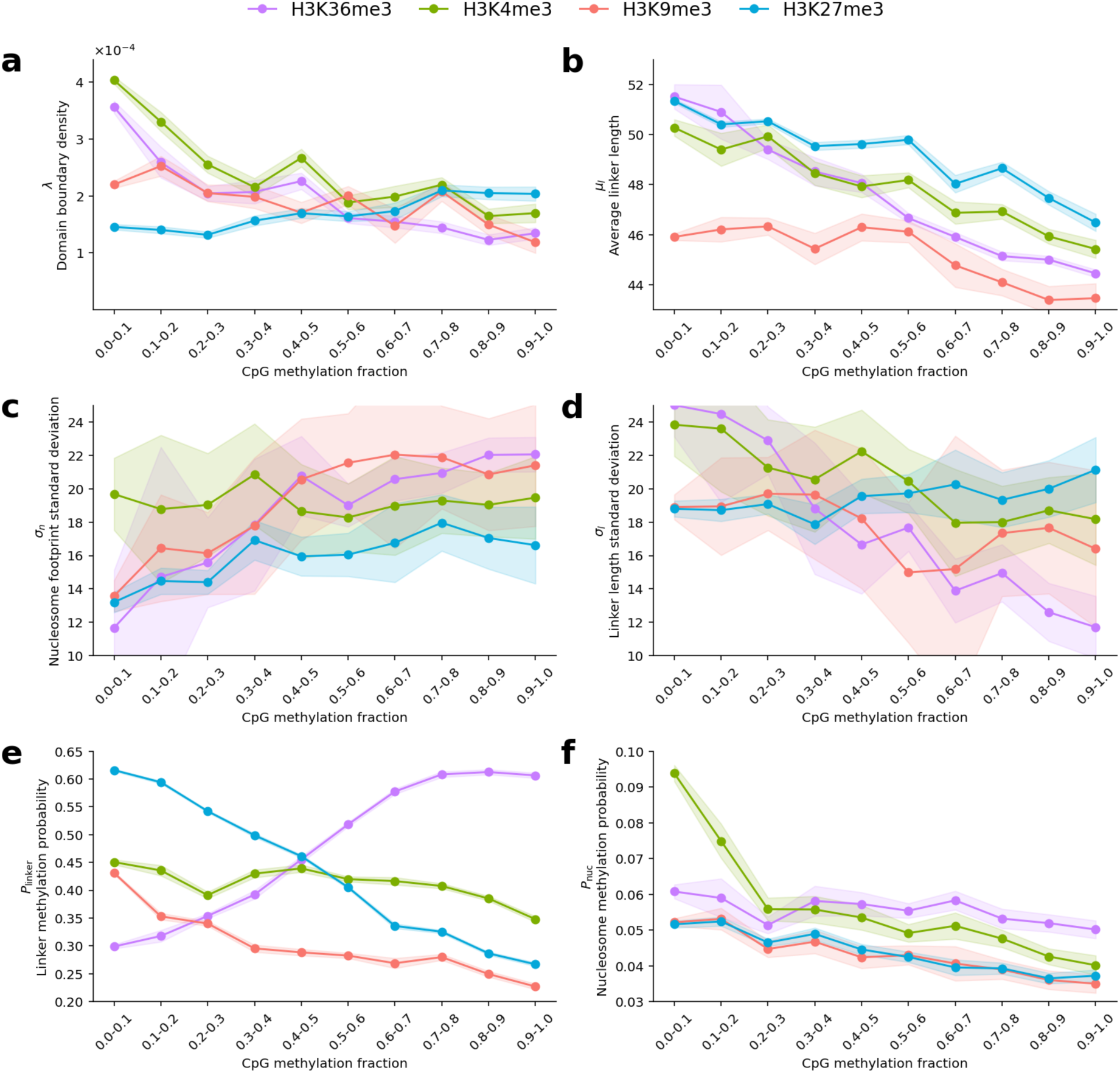
Model parameters inferred from single-molecule long reads across epigenetic states in K562 cells. Each panel shows a different inferred parameter of the nucleosome–linker model (y-axis) as a function of CpG methylation fraction (x-axis) for four histone modifications (H3K36me3, H3K4me3, H3K9me3, H3K27me3 – colour). Shaded regions in all panels represent estimated 95% confidence intervals of the inferred parameters, calculated from the covariance matrix obtained from the Jacobian of the least-squares fit. **a.** Domain boundary density λ, representing the probability per base that a read transitions out of a histone-modified region. Larger λ corresponds to shorter continuous modification regions. **b.** Average linker length μ_l_ corresponding to the mean of the gamma distribution underlying L_linker_. **c.** Nucleosome footprint standard deviation σ_n_, corresponding to the standard deviation of the gamma distribution underlying L_nuc_. **d.** Linker length standard deviation σ_l_, corresponding to the standard deviation of the gamma distribution underlying L_linker_. **e.** Linker methylation probability, P_linker_. **f.** Nucleosome methylation probability, P_nuc_.

**Supplementary Figure 8.**
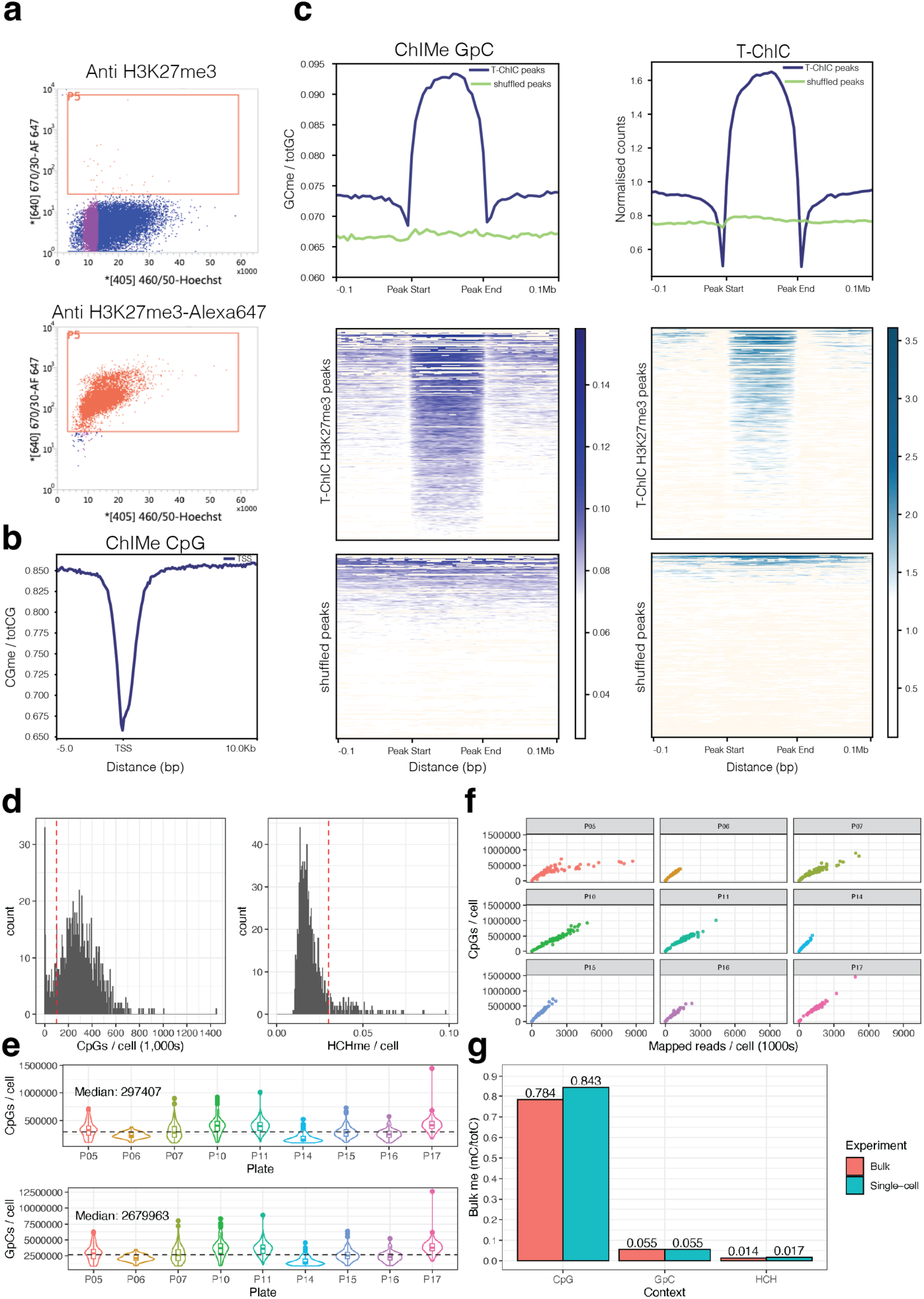
Global performance metrics for single-cell ChIMe-seq after processing with scBS-seq. **a.** FACS plots showing Hoechst 34580 signal Vs Alexa-647 signal of day 6 mouse gastruloid cells processed with ChIMe-seq with or without an Alexa-647 conjugated H3K27me3 antibody. **b.** Average CpG methylation around all transcription start sites (TSSs) across all day 5 mouse gastruloid cells processed with ChIMe-seq. **c.** Average H3K27me3 signal (GpC methylation – ChIMe-seq, normalized counts – T-ChIC) around all H3K27me3 peaks, called from T-ChIC, across all day 5 mouse gastruloid cells, or across shuffled peaks. Heatmaps show signal across individual peaks. **d.** Histograms of i) number of CpGs detected per cell in ChIMe-seq, ii) HCH methylation fraction per cell in ChIMe-seq, red lines indicate the thresholds cells are filtered on. **e.** Violin plots of CpGs or GpCs detected per cell in ChIMe-seq, separated by plate. Black line indicates median across all cells. **f.** Oversequencing plots for ChIMe-seq showing mapped read counts Vs CpGs detected per cell, separated by plate. **g.** Barplot showing mean methylation values across contexts for: red, bulk processing of gastruloid sample and blue, single-cell processing of all, post-filtering, single-cells collected from the same ChIMe-seq preparation.

**Supplementary Figure 9.**
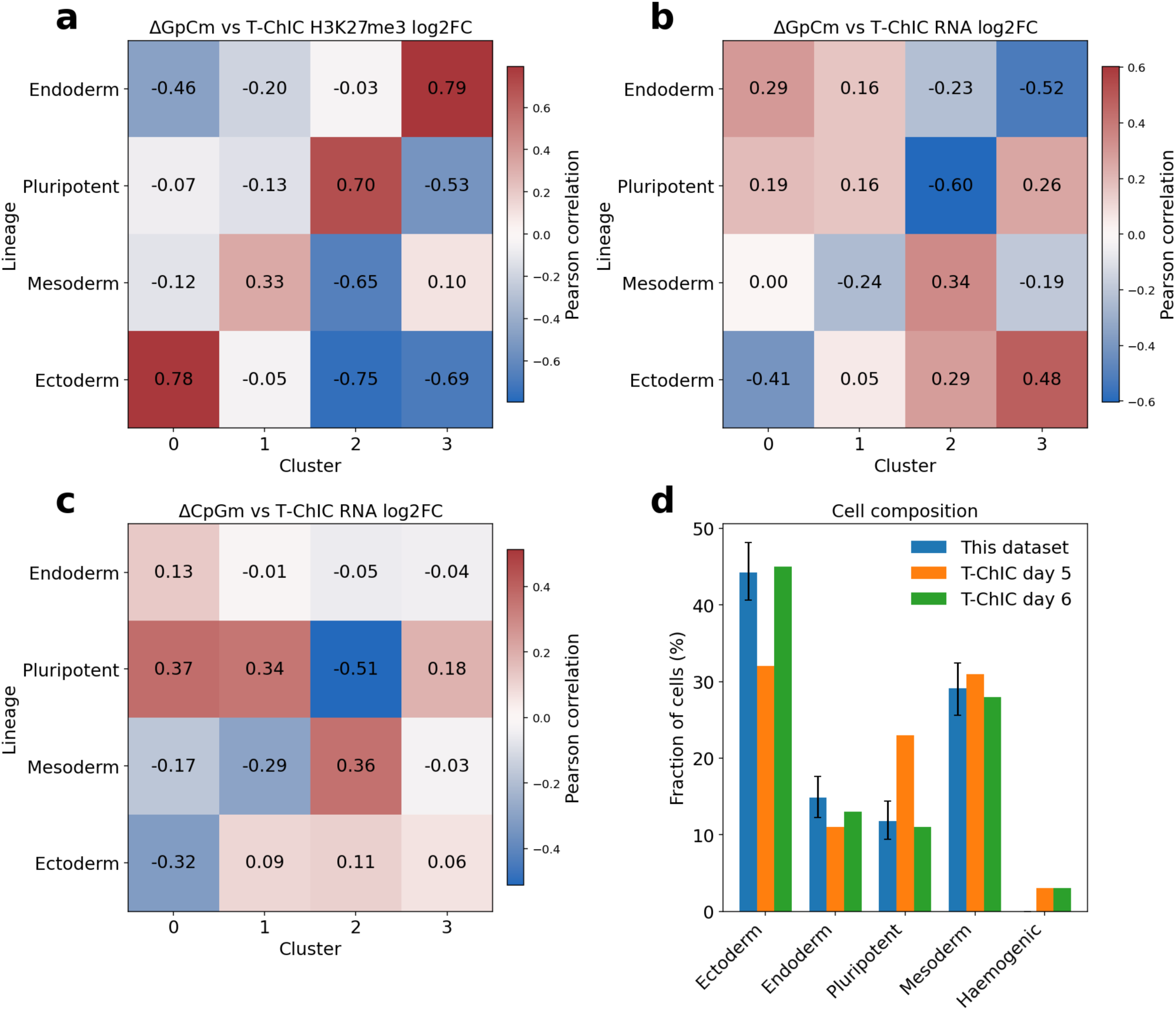
Annotation and validation of ChIMe-derived cell clusters using transcriptomic and chromatin datasets. **a.** Comparison between differential GpC methylation, measured by ChIMe-seq and lineage-specific H3K27me3 signal enrichment from T-ChIC, for each cluster-lineage pair. Comparisons made across cluster specific differentially methylated regions (DMRs) identified in ChIMe-seq GpC methylation. ΔGpCme; the difference in mean DMR H3K27me3 directed GpC methylation fraction between ChIMe-seq clusters. T-ChIC H3K27me3 log2 fold change; the difference in ChIC read detection across ChIMe-seq DMRs. **b.** Correlation between differential GpC methylation, measured by ChIMe-seq, and lineage-specific differential gene expression, derived from T-ChIC RNA-seq, for each cluster-lineage pair. Comparisons made across cluster specific differentially methylated regions (DMRs) identified in ChIMe-seq GpC methylation. ΔGpCme; as in **a**. T-ChIC log2 fold change; the difference in RNA expression of DMR linked genes between T-ChIC lineages. As H3K27me3 is associated with transcriptional repression, matching lineage–cluster pairs are expected to exhibit negative correlations. **c.** Correlation between differential promoter CpG methylation, measured by ChIMe-seq, and lineage-specific differential gene expression, derived from T-ChIC RNA-seq, for each cluster-lineage pair. Calculated as in panel **b**, but using ChIMe-seq CpG methylation derived DMRs and ΔCpGme for ChIMe-seq cluster comparison. As promoter DNA methylation is generally associated with transcriptional repression, matching lineage–cluster pairs are expected to exhibit negative correlations. **d.** Comparison of lineage representation proportions inferred from ChIMe-seq clusters with RNA based T-ChIC reported lineage proportions from day 5 and day 6 gastruloids.

**Supplementary Figure 10.**
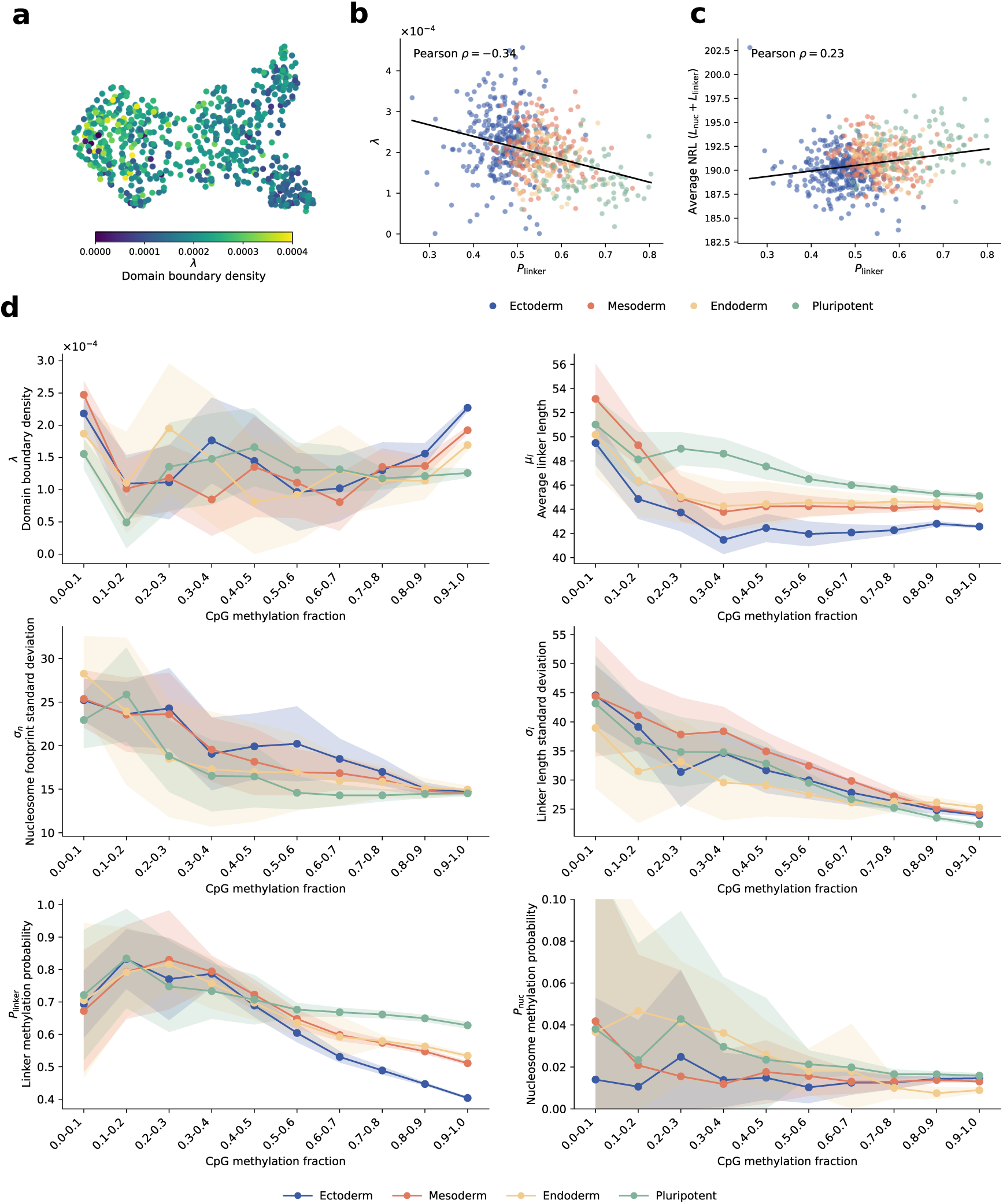
Model parameters inferred across DNA methylation states in day 5 mouse gastruloids. **a.** Integrated CpG–GpC UMAP embedding coloured by the fitted domain boundary density λ, representing the probability per base that of transitioning out of a histone-modified region. **b.** Single-cell relationship between linker methylation probability P_linker_ and domain boundary density λ. Points represent individual cells coloured by gastruloid lineage, with the black line indicating the linear regression fit and the Pearson correlation coefficient shown. **c.** Single-cell relationship between linker methylation probability P_linker_ and average nucleosome repeat length (NRL), displayed as in **b**. **d.** Model parameters as a function of CpG methylation fraction for the four gastruloid lineages. Lines and points represent mean parameter estimates across 100 bootstrap refits, generated by resampling cells and repeating CpG methylation stratification, conditional GpC methylation probability calculation and model fitting. Shaded regions indicate ±1 s.d. across bootstrap refits.

**Supplementary Table 1.** Comparison of Fraction of Signal in Peaks for ChIMe-seq and published immuno-targeting histone modification profiling methods. Values represent the fraction of individual GpC methylation calls found within peaks called from ENCODE ChIP-seq data for ChIMe-seq and the self-reported fraction of individual mapped reads found within target peaks in published datasets. Missing values are not reported in the listed study.

| **Publication** | **Method** | **H3K27me3** | **H3K9me3** | **H3K4me3** | **H3K36me3** | **Effector Enzyme** | **Cells** |
|---|---|---|---|---|---|---|---|
| This publication | ChIMe-seq | 0.77 | 0.75 | 0.12 | 0.77 | M.CviPI | K562 |
| Kaya-Okur et al<br>PMID: 31036827 | CUT&Tag (2019) | 0.34 |  |  |  | Tn5 | K562 |
| Harada et al<br>PMID: 30532068 | ChIL-seq (2019) | 0.1 |  | 0.4 |  | Tn5 | C2C12 |
| Ku et al<br>PMID: 30923384 | scChIC-seq (2019) | 0.47 |  | 0.5 |  | MNase | WBCs |
| Li et al<br>PMID: 35028637 | CUT&Tag-BS (2021) | 0.61 | 0.19-0.26 |  |  | Tn5 | mESCs |
| Meers et al<br>PMID: 36316484 | MulTI-Tag (2022) | 0.068 |  |  | 0.178 | Tn5 | H1 hESC |
| Zeller et al<br>PMID: 36539617 | sortChIC (2023) | 0.89 | 0.73 | 0.71 |  | MNase | K562 |
| Bartosovic et al<br>PMID: 36536148 | Nano-CT (2023) | 0.39 |  |  |  | Tn5 | Mouse Brain |
| Li et al<br>PMID: 39375575 | scNanoSeq-CUT&Tag (2024) | ~0.32 | ~0.26 | ~0.81 | ~0.65 | Tn5 | K562, 293T, GM12878, HG002, H9 and HFF1 |
| Geisenberger et al<br>PMID: 40999097 | scEpi<sup>2</sup>-seq (2025) | 0.88 | 0.72 |  | 0.83 | MNase | K562 |

