## Supplementary Note for "ChIMe-seq enables multimodal epigenetic profiling in single molecules and single cells"

### Supplementary Information

Joe Verity-Legg, Euan Joly-Smith, Roy Straver, Soufiane Mourragui, Stephen Clark, Vivek Bhardwaj, Marloes Blotenburg, Francesca Mattiroli, Sito Torres-Garcia, Wolf Reik, Jeroen de Ridder, Peter Zeller, Alexander van Oudenaarden

**Contents**

Sequences 2
A. Nucleotide sequence 2
B. Amino Acid sequence 2
C. Nucleotide sequence — in sections 2
ORF 2
Protein A 2
6x HA tags 3
6 His tags 3
Linker 3
M.CviPI 3

Nucleosome–linker model 4

I. Description of the model in histone-modified domains 4

II. Transition matrix representation 5

III. Conditional linker-state probabilities 6

IV. Incorporating GpC methylation 8

V. Histone-modification domain decay 9

VI. Extension to two alleles 11

VII. Model fitting procedure 13
A. Fitting the model from the first oscillation peak 13
B. Direct propagation of state probabilities 13
C. Model fitting and parameter estimation 14

References 14

#### A. Nucleotide sequence

ATGATTACGCCAAGCTTAAAAGATGACCCAAGCCAAAGTGCTAACCTATTGTCAGAAGCTAAAAAGTT
AAATGAATCTCAAGCACCGAAAGCGGATAACAAATTCAACAAAGAACAACAAAATGCTTTCTATGAAA
TCTTACATTTACCTAACTTAAACGAAGAACAACGCAATGGTTTCATCCAAAGCCTAAAAGATGACCCA
AGCCAAAGCGCTAACCTTTTAGCAGAAGCTAAAAAGCTAAATGATGCTCAAGCACCAAAAGCTGACAA
CAAATTCAACAAAGAACAACAAAATGCTTTCTATGAAATTTTACATTTACCTAACTTAACTGAAGAAC
AACGTAACGGCTTCATCCAAAGCCTTAAAGACGATCCTTCAGTGAGCAAAGAAATTTTAGCAGAAGCT
AAAAAGCTAAACGATGCTCAAGCACCAAAAGATGACTATCCCTATGACGTCCCTGATTACGCGTACCC
GTACGATGTCCCCGATTATGCCTACCCTTATGACGTGCCCGATTACGCCCACCACCACCACCACCATG
GGCCAGGTTCGGGTGGGGGTGGTGGCGGTAGTGGTGGCTCAGGAGGTGGCCTCGGTTCGACAACCTT
GAAAGCCCTCGAATTATTCGCCGGAATTGCGGGAATAACGCATGGTCTTCGTGGATTCGTAGAACCCG
TGGCTTTCGTCGAGATAAACAAAGATGCTCAAGAATTTCTCTCTACAAAGTTCCCGGATAAACCGGTG
TTCGATGATGTGACGAAATTTTCAAAACGTGATTTTGACGAGCCGATCGATATGATTACCGGAGGATT
CCCTTGCACCGGTTTTAGCATAGCAGGTAAACGAAATGGTTTCGAACACGCAGAATCCGGCTTATTCG
GAGAAGTCGTGCGCATAACGAAAGAATACATGCCAAAGATGGTCTTTTTAGAAAACTCTGGCATGTTG
AGTCACAAGTATAACTTGGACATCGTCATCAGATCTATGGATTCTCTAGGCTATGATTGTCGATGGGT
TACTTTGCGAGCCACAGTCGTGGGAGCTTTACACACGAGACATCGTTGGTTCTGTTTGTGCACACGTA
AAGATCATATTCGCGAAACGCTCATTTGCGATCGAGAAGTCACTAAGTTCGACTGGGAAAATGATAGA
CCTCCTATACAGGTAGACTCACGAAGCTATGAAAATAGTCGTCTCGTGAGATTTGCCGGTTATTCCGT
CGTTCCCGATCAAATTCGATATGCATTCACAGGTCTCTACACCGGAAATTTCTCACCATCGTTCTCGAA
GACGCTCGTACCAGGATCATTAGAAGGAAGTATTTGTTTCAATGAAGACAAAATAACGAACGGATATT
ACAAAGATGGTGTATATTATGAATTCGTTCGCACGGAGACACACAGAGAACCCGTGAACATACTTCTG
ACTCCGAGAGAAATACCGAATAAACATAACGGGAAAAAACTGCTCACTTTACCAGTGACGAAAAGATA
TTGGTGTACTCCTTGTGCTTCATACGGAAAAGGAACCGCAGGAGGCAGAGTATTAACAGACCGTTCGA
GTCATTCTCTTCCAACTCAAGTGAAATTTTCTCCCGAAGGAGAGGACGGAAAACATCTATCTGGAAAG
TTTTGTGCATGGCTCATGGGATATGACAAAGAATATTTAGGAAATTTGTTAGAATATTGA

ORF - pA domain - HAx6 (6 HA tags) domain - His tag - Linker - M.CviPI

#### B. Amino Acid sequence

MITPSLKDDPSQSANLLSEAKKLNESQAPKADNKFNKEQQNAFYEILHLPNLNEEQRNGFIQSLKDDPSQSAN
LLAEAKKLNDAQAPKADNKFNKEQQNAFYEILHLPNLTEEQRNGFIQSLKDDPSVSKEILAEAKKLNDAQAP
KDDYPYDVPDYAYPYDVPDYAYPYDVPDYAHHHHHHGPGSGGGGGGSGGSGGGLGSTTLKALELFAGIAG
ITHGLRGFVEPVAFVEINKDAQEFLSTKFPDKPVFDDVTKFSKRDFDEPIDMITGGFPCTGFSIAGKRNGFEH
AESGLFGEVVRITKEYMPKMVFLENSGMLSHKYNLDIVIRSMDSLGYDCRWVTLRATVVGALHTRHRWFCL
CTRKDHIRETLICDREVTKFDWENDRPPIQVDSRSYENSRLVRFAGYSVVPDQIRYAFTGLYTGNFSPSFSKT
LVPGSLEGSICFNEDKITNGYYKDGVYYEFVRTETHREPVNILLTPREIPNKHNGKKLLTLPVTKRYWCTPC
ASYGKGTAGGRVLTDRSSHSLPTQVKFSPEGEDGKHLSGKFCAWLMGYDKEYLGNLLEY*

#### C. Nucleotide sequence — in sections

*ORF*

ATGATTACGCCAAGCTTAAAAGATGACCCAAGCCAAAGTGCTAACCTATTGTCAGAAGCTAAAAAGTT
AAATGAATCTCAAGCACCGAAAGCGGATAACAAATTCAACAAAGAACAACAAAATGCTTTCTATGAAA
TCTTACATTTACCTAACTTAAACGAAGAACAACGCAATGGTTTCATCCAAAGCCTAAAAGATGACCCA
AGCCAAAGC

*Protein A*

GCTAACCTTTTAGCAGAAGCTAAAAAGCTAAATGATGCTCAAGCACCAAAAGCTGACAACAAATTCAA
CAAAGAACAACAAAATGCTTTCTATGAAATTTTACATTTACCTAACTTAACTGAAGAACAACGTAACG
GCTTCATCCAAAGCCTTAAAGACGATCCTTCAGTGAGCAAAGAAATTTTAGCAGAAGCTAAAAAGCTA
AACGATGCTCAAGCACCAAAAGATGAC

*6x HA tags*

TATCCCTATGACGTCCCTGATTACGCGTACCCGTACGATGTCCCCGATTATGCCTACCCTTATGACGT
GCCCGATTACGCC

*6 His tags*

CACCACCACCACCACCAT

*Linker*

GGGCCAGGTTCGGGTGGGGGTGGTGGCGGTAGTGGTGGCTCAGGAGGTGGCCTCGGTTCGACA

*M.CviPI*

ACCTTGAAAGCCCTCGAATTATTCGCCGGAATTGCGGGAATAACGCATGGTCTTCGTGGATTCGTAGA
ACCCGTGGCTTTCGTCGAGATAAACAAAGATGCTCAAGAATTTCTCTCTACAAAGTTCCCGGATAAAC
CGGTGTTCGATGATGTGACGAAATTTTCAAAACGTGATTTTGACGAGCCGATCGATATGATTACCGGA
GGATTCCCTTGCACCGGTTTTAGCATAGCAGGTAAACGAAATGGTTTCGAACACGCAGAATCCGGCTT
ATTCGGAGAAGTCGTGCGCATAACGAAAGAATACATGCCAAAGATGGTCTTTTTAGAAAACTCTGGCA
TGTTGAGTCACAAGTATAACTTGGACATCGTCATCAGATCTATGGATTCTCTAGGCTATGATTGTCGA
TGGGTTACTTTGCGAGCCACAGTCGTGGGAGCTTTACACACGAGACATCGTTGGTTCTGTTTGTGCAC
ACGTAAAGATCATATTCGCGAAACGCTCATTTGCGATCGAGAAGTCACTAAGTTCGACTGGGAAAATG
ATAGACCTCCTATACAGGTAGACTCACGAAGCTATGAAAATAGTCGTCTCGTGAGATTTGCCGGTTAT
TCCGTCGTTCCCGATCAAATTCGATATGCATTCACAGGTCTCTACACCGGAAATTTCTCACCATCGTTC
TCGAAGACGCTCGTACCAGGATCATTAGAAGGAAGTATTTGTTTCAATGAAGACAAAATAACGAACGG
ATATTACAAAGATGGTGTATATTATGAATTCGTTCGCACGGAGACACACAGAGAACCCGTGAACATAC
TTCTGACTCCGAGAGAAATACCGAATAAACATAACGGGAAAAAACTGCTCACTTTACCAGTGACGAAA
AGATATTGGTGTACTCCTTGTGCTTCATACGGAAAAGGAACCGCAGGAGGCAGAGTATTAACAGACCG
TTCGAGTCATTCTCTTCCAACTCAAGTGAAATTTTCTCCCGAAGGAGAGGACGGAAAACATCTATCTG
GAAAGTTTTGTGCATGGCTCATGGGATATGACAAAGAATATTTAGGAAATTTGTTAGAATATTGA

### Nucleosome–linker model

This Supplementary Note provides the mathematical derivations of the nucleosome–linker model used to fit the pairwise GpC methylation probabilities. We first describe nucleosome organization within a histone-modified domain and define the distributions of nucleosome-protected and linker lengths (Sec. I). We then construct the transition matrix (Sec. II) and use it to derive the conditional probability that a nucleotide is in a linker state given that another nucleotide a distance $x$ away is also in a linker state (Sec. III). We next relate this latent-state conditional probability to the experimentally measured pairwise conditional GpC methylation probability (Sec. IV) and extend the model to account for the finite extent of histone-modified domains and the possibility of crossing a domain boundary (Sec. V). Finally, for fitting the single-cell oscillations, we extend the model to account for contributions from the two alleles and differences in their nucleosome–linker organization (Sec. VI).

#### I. Description of the model in histone-modified domains

We model DNA as a one-dimensional lattice in which each lattice point corresponds to a nucleotide. Each nucleotide can exist in one of two states. If a nucleotide is wrapped around a nucleosome, we say that it is in state $n$. If a nucleotide is located within a linker region between nucleosomes, we say that it is in state $l$ (Fig. S1a). Consecutive nucleotides in state $n$ therefore form nucleosome-protected regions, whereas consecutive nucleotides in state $l$ form linker regions. We allow the lengths of both regions to vary.

We first consider this nucleosome–linker organization within a histone-modified domain. We define a histone-modified domain as a genomic region in which artificial GpC methylation can be deposited. GpCs outside histone-modified domains are assumed to be unmethylated in the model.

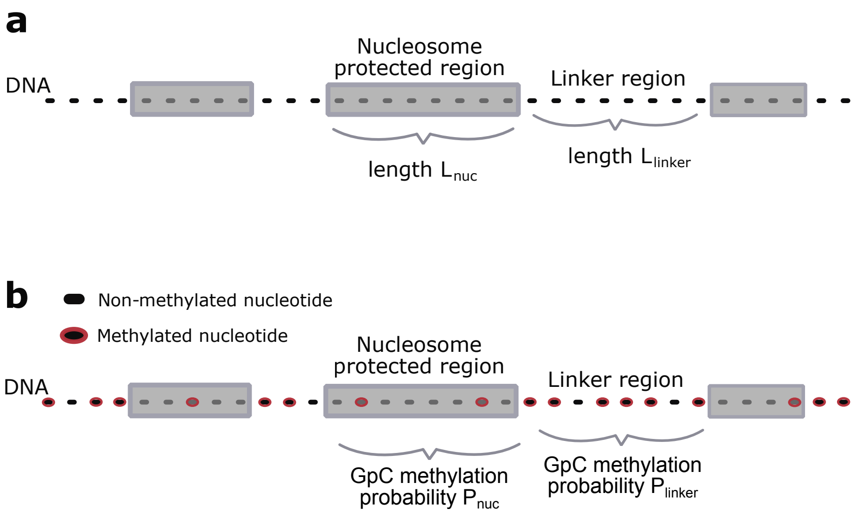


Figure S1. **A model for nucleosome spacing along DNA. a)** Nucleosomes are distributed along the DNA, generating nucleosome-protected and linker regions across the nucleotide lattice (black dots). The lengths of nucleosome-protected and linker regions are variable and are each modelled by a Gamma distribution (Eq. (1)). Throughout this work, the mean nucleosome-protected region length is fixed at 147 bp. **b)** Whether a nucleotide is within a nucleosome-protected or linker region is not directly observed. Instead, the reference sequence identifies GpC sites, and the base-calling pipeline determines whether the cytosine at each GpC site is methylated. We define $P_{\text{linker}}$ and $P_{\text{nuc}}$ as the probabilities that a known GpC site is measured as methylated when it lies within a linker or nucleosome-protected region, respectively.

For a randomly selected nucleosome-protected region, its length $L_{\text{nuc}}$ is a random variable with probability distribution $D_{\text{nuc}}$:

$$L_{\text{nuc}} \sim D_{\text{nuc}}(x),$$

where $D_{\text{nuc}}(x)$ describes the probability of a nucleosome-protected region having length $x$. Similarly, the length $L_{\text{linker}}$ of a randomly selected linker region is a random variable with probability distribution $D_{\text{linker}}$:

$$L_{\text{linker}} \sim D_{\text{linker}}(x),$$

where $D_{\rm linker}(x)$ describes the probability of a linker region having length $x$.

The derivations below hold for arbitrary choices of $D_{\rm nuc}(x)$ and $D_{\rm linker}(x)$. Here, we model both using Gamma distributions:

$$\begin{aligned} D_{\rm nuc}(x) &= \frac{1}{\Gamma(\alpha_n)\theta_n^{\alpha_n}} x^{\alpha_n-1} e^{-x/\theta_n}, \qquad \alpha_n = \frac{\mu_n^2}{\sigma_n^2}, \qquad \theta_n = \frac{\sigma_n^2}{\mu_n}, \\ D_{\rm linker}(x) &= \frac{1}{\Gamma(\alpha_l)\theta_l^{\alpha_l}} x^{\alpha_l-1} e^{-x/\theta_l}, \qquad \alpha_l = \frac{\mu_l^2}{\sigma_l^2}, \qquad \theta_l = \frac{\sigma_l^2}{\mu_l}, \end{aligned} \tag{1}$$

where $\mu_n$ and $\sigma_n$ are the mean and standard deviation of nucleosome-protected region lengths, and $\mu_l$ and $\sigma_l$ are the mean and standard deviation of linker region lengths.

Eq. (1) describes the underlying nucleosome organization in terms of the nucleotide states $n$ and $l$. These states are defined at every nucleotide position but are not measured directly. Instead, the reference sequence identifies the positions of GpC sites, and at each such position we measure whether the cytosine is methylated. The probability that a GpC site is measured as methylated depends on whether the corresponding nucleotide position is in state $n$ or $l$. GpC sites within nucleosome-protected regions are less accessible to artificial methylation than those within linker regions. We therefore define

$$\begin{aligned} P_{\rm nuc} &= P(\text{methylated} \mid \text{GpC site}, n), \\ P_{\rm linker} &= P(\text{methylated} \mid \text{GpC site}, l). \end{aligned} \tag{2}$$

Thus, $P_{\rm nuc}$ and $P_{\rm linker}$ describe methylation probabilities conditional on the position being a known GpC site. They do not include the probability that a nucleotide occurs in a GpC context (Fig. S1b).

The model therefore contains six parameters:

$$\mu_n,\ \mu_l,\ \sigma_n,\ \sigma_l,\ P_{\rm nuc},\ P_{\rm linker}.$$

We constrain the model by fixing the mean nucleosome-protected region length to 147 bp, such that $\mu_n = 147$ bp. This leaves five free parameters to be inferred from the data.

#### II. Transition matrix representation

DNA consists of alternating nucleosome-protected and linker regions whose lengths are drawn from the distributions $D_{\rm nuc}$ and $D_{\rm linker}$. To represent these length distributions, we expand the two states $n$ and $l$ into substates that specify the position of a nucleotide within its current nucleosome-protected or linker region.

In particular, each nucleotide can take one of $N = N_n + N_l$ substates. If a site is in state $k \le N_n$, then it is located in a nucleosome protected region, at position $k$ starting from left side of the region. If a site is in substate $N_n < k$, then it is located in a linker region, a distance $k - N_n$ from the nearest nucleosome on the left of the position. $N_n$ and $N_l$ then effectively set the maximum possible lengths of the nucleosome protected and linker regions respectively, see Fig. S2

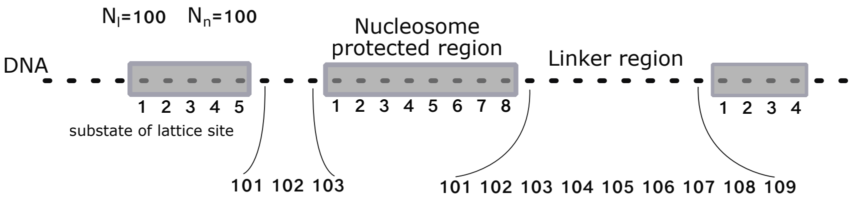


Figure S2. **Illustration of the substate definitions for each lattice site.** Each lattice site (i.e., nucleotide) can take one of $N = N_n + N_l$ substates. If a site is in state $k \le N_n$, then it is located in a nucleosome protected region, at position $k$ starting from left side of the region. If a site is in substate $N_n < k$, then it is located in a linker region, a distance $k - N_n$ from the nearest nucleosome on the left of the position. $N_n$ and $N_l$ then effectively set the maximum possible lengths of the nucleosome protected and linker regions respectively. We can set these to be large in the regime where the probability of finding such lengths ($P_n(N_n)$ and $P_l(N_l)$ in Eq. (1)) are approximately zero. Shown here is an example where $N_n = 100$ and $N_l = 100$. Each lattice site is a black dot, with numbers underneath correspond to substate indices for each site.

The dimensions of the transfer matrix will be set by the number of substates $N = N_n + N_l$. The number of substates thus needs to be finite in order for the transfer matrix approach to be computationally possible. In addition, the

Gamma distributions in Eq. (1) are continuous, whereas the region lengths on the nucleotide lattice take integer values. We thus set $N_n$ to be the smallest $A$ such that $\sum_{k=1}^{A} D_{\text{nuc}}(k) > 0.99$ in order for the substates to span 99% of the distibution. $N_l$ is chosen similarly but with $D_{\text{linker}}$. The resulting probabilities are renormalized over the represented lengths to have total cumulative probability of 1:

$$P_n(r) = \frac{D_{\text{nuc}}(r)}{\sum_{k=1}^{N_n} D_{\text{nuc}}(k)}, \qquad r = 1, \ldots, N_n, \qquad P_l(r) = \frac{D_{\text{linker}}(r)}{\sum_{k=1}^{N_l} D_{\text{linker}}(k)}, \qquad r = 1, \ldots, N_l.$$

Let

$$C_n(r) = \sum_{k=1}^{r} P_n(k), \quad C_l(r) = \sum_{k=1}^{r} P_l(k)$$

denote the corresponding cumulative distributions. Given that a nucleosome-protected region has reached length $r$, the conditional probability that it stops the current position and becomes a linker state at the next position is (which follows from Baye's rule)

$$f_n(r) = P(L_{\text{nuc}} = r \mid L_{\text{nuc}} \geq r) = \frac{P_n(r)}{1 - C_n(r-1)}. \tag{3}$$

Similarly,

$$f_l(r) = P(L_{\text{linker}} = r \mid L_{\text{linker}} \geq r) = \frac{P_l(r)}{1 - C_l(r-1)}. \tag{4}$$

We can describe movement along the nucleotide lattice using a transition matrix $T$, whose elements are

$$T_{ij} = P(S_{x+1} = j \mid S_x = i),$$

where $S_x$ denotes the substate at nucleotide position $x$. For a nucleosome substate $k < N_r$ there are two possibilities if we move one nucleotide to the right: either the neighbouring state will be $k+1 \leq N_r$ thus remaining in a nucleosome-protected region or it will switch to linker and be in substate $N_n + 1$. These two transition probabilities are given by

$$T_{k,k+1} = 1 - f_n(k) \text{ and } T_{k,N_n+1} = f_n(k),$$

respectively. When $k = N_n$ there is just one possibility: the next nucleotide will switch to linker and thus be in the $N_n + 1$ substate. Thus

$$T_{N_n,N_n+1} = 1.$$

Similarly, for a linker substate $N_n + k$ where now $k < N_l$, we have

$$T_{N_n+k,N_n+k+1} = 1 - f_l(k) \text{ and } T_{N_n+k,1} = f_l(k),$$

and additionally

$$T_{N_n+N_l,1} = 1.$$

All other elements of $T$ are zero. Each row therefore sums to one.

##### III. Conditional linker-state probabilities

We next calculate the probability that a nucleotide at distance $x$ is in a linker state given that the nucleotide at position 0 is in a linker state.

Let

$$\boldsymbol{\pi}^{\mathsf{T}} = (\pi_1, \ldots, \pi_{N_n}, \pi_{N_n+1}, \ldots, \pi_{N_n+N_l})$$

denote the distribution over substates, such that $\pi_i$ is the probability that a randomly selected nucleotide is in substate $i$.

More generally, let $\boldsymbol{\pi}^{\mathsf{T}}(x)$ denote the substate distribution at nucleotide position $x$. Multiplication by the transition matrix propagates this distribution by one nucleotide:

$$\boldsymbol{\pi}^{\mathsf{T}}(x+1) = \boldsymbol{\pi}^{\mathsf{T}}(x)T.$$

We assume that the nucleosome-linker process is stationary, such that the substate distribution is independent of position. Therefore,

$$\boldsymbol{\pi}^{\mathsf{T}}T = \boldsymbol{\pi}^{\mathsf{T}}, \qquad \sum_i \pi_i = 1. \tag{5}$$

Consider the linker substates $N_n+1, N_n+2, \ldots, N_n+N_l$. In the transition matrix $T$, the only transition into substate $N_n+k$, where $k>1$, is from the preceding substate $N_n+k-1$. The transition probability is

$$T_{N_n+k-1,N_n+k} = 1 - f_l(k-1).$$

Thus at stationarity,

$$\pi_{N_n+k} = \pi_{N_n+k-1}T_{N_n+k-1,N_n+k} = \pi_{N_n+k-1}\left[1-f_l(k-1)\right].$$

It follows that

$$\pi_{N_n+k} = \pi_{N_n+1}\prod_{i=1}^{k-1}\left[1-f_l(i)\right]. \tag{6}$$

The product in Eq. (6) is the probability that a linker region does not stop and switch to a nucleosome region before reaching substate $N_n+k$. Thus it is equal to the probability that its length is at least $k$:

$$P(L_{\text{linker}} \geq k) = \prod_{i=1}^{k-1}\left[1-f_l(i)\right].$$

Thus,

$$\pi_{N_n+k} = \pi_{N_n+1}P(L_{\text{linker}} \geq k). \tag{7}$$

Using Eq. (7) and conditioning position 0 on being in a linker region gives

$$P(S_0 = N_n+k \mid S_0 \in l) = \frac{P(L_{\text{linker}} \geq k)}{\displaystyle\sum_{i=1}^{N_l} P(L_{\text{linker}} \geq i)} \tag{8}$$

where $S_x$ is defined as the substate of position $x$ on the lattice, and $S_0 \in l$ means the substate of position 0 is in linker state.

We define the following vector

$$\mathbf{q}^{\mathsf{T}} = (0, \ldots, 0, q_1, \ldots, q_{N_l}),$$

where $q_k := P(S_0 = N_n+k \mid S_0 \in l)$. Thus $\mathbf{q}^{\mathsf{T}}$ consists of the probability distribution of substates for position 0 of the DNA conditioned on that position 0 is in a linker state. After moving $x$ nucleotides along the DNA, the conditional distribution over substates $\mathbf{p}_x$ is

$$\mathbf{p}_x^{\mathsf{T}} = \mathbf{q}^{\mathsf{T}}T^x. \tag{9}$$

To obtain the probability that position $x$ is in a linker state, we define the indicator vector

$$\mathbf{g}_l = (0, \ldots, 0, 1, \ldots, 1)^{\mathsf{T}},$$

where the final $N_l$ entries correspond to linker substates. The probability that position $x$ is in a linker state given that position 0 is in a linker state is then

$$\rho_l(x \mid 0) = P(S_x \in l \mid S_0 \in l) = \mathbf{q}^{\mathsf{T}}T^x\mathbf{g}_l. \tag{10}$$

#### IV. Incorporating GpC methylation

The model above describes the underlying nucleosome organization in terms of the latent nucleotide states $n$ and $l$. These states are not measured directly. Instead, at known GpC sites we measure whether the cytosine is methylated. We denote the event that the GpC site at position $x$ is measured as methylated by $M_x$.

As defined in Eq. (2), the probability of methylation depends on the underlying nucleotide state:

$$P(M_x \mid S_x = n) = P_{\text{nuc}},$$
$$P(M_x \mid S_x = l) = P_{\text{linker}}.$$

We aim to find the probability that a GpC site at position $x$ is methylated conditional on a GpC site at position 0 also being methylated,

$$P(M_x \mid M_0).$$

Conditioning on the underlying state at position 0 gives

$$P(M_x \mid M_0) = P(M_x \mid S_0 = l, M_0)P(S_0 = l \mid M_0) + P(M_x \mid S_0 = n, M_0)P(S_0 = n \mid M_0). \tag{11}$$

Once the underlying state at position 0 is specified, the methylation measurement at position 0 provides no additional information about the state at position $x$. Therefore,

$$P(M_x \mid S_0 = l, M_0) = P(M_x \mid S_0 = l),$$
$$P(M_x \mid S_0 = n, M_0) = P(M_x \mid S_0 = n).$$

For a stationary sequence of alternating nucleosome-protected and linker regions, the probabilities that a randomly selected nucleotide lies in either state are

$$\pi_n = P(S_x = n) = \frac{\mu_n}{\mu_n + \mu_l},$$
$$\pi_l = P(S_x = l) = \frac{\mu_l}{\mu_n + \mu_l}.$$

The mean methylation probability of a known GpC site is therefore

$$\rho_M = P(M_x) = P_{\text{nuc}}\pi_n + P_{\text{linker}}\pi_l = \frac{P_{\text{nuc}}\mu_n + P_{\text{linker}}\mu_l}{\mu_n + \mu_l}. \tag{12}$$

Bayes' rule gives the probability that a methylated GpC site lies within either state:

$$P(S_0 = n \mid M_0) = \frac{P_{\text{nuc}}\mu_n}{P_{\text{nuc}}\mu_n + P_{\text{linker}}\mu_l}, \tag{13}$$

$$P(S_0 = l \mid M_0) = \frac{P_{\text{linker}}\mu_l}{P_{\text{nuc}}\mu_n + P_{\text{linker}}\mu_l}. \tag{14}$$

We now consider $P(M_x \mid S_0 = l)$ and write it in terms of the conditional linker probability expression from Eq. (10):

$$P(M_x \mid S_0 = l) = P_{\text{linker}}\rho_l(x \mid 0) + P_{\text{nuc}}\left[1 - \rho_l(x \mid 0)\right], \tag{15}$$

where we used the fact that in this model the methylation probability at a particular position is set by the underlying state of that position (and so conditioning on additional states of other positions does not add any information, i.e., $P(M_x \mid S_x = n, S_0 = l) = P(M_x \mid S_x = n) = P_{\text{nuc}}$ for example.

To obtain the corresponding expression conditional on position 0 being nucleosome-protected, we first note that

$$1 - \rho_l(x \mid 0) = P(S_x = n \mid S_0 = l).$$

Stationarity and spatial symmetry of the alternating-region model imply that the joint probability of observing a linker state at position 0 and a nucleosome-protected state at position $x$ is equal to the joint probability of observing the reverse configuration:

$$P(S_0 = l, S_x = n) = P(S_0 = n, S_x = l).$$

Expanding each joint probability using $P(A, B) = P(A)P(B \mid A)$ gives

$$P(S_0 = l, S_x = n) = P(S_0 = l)P(S_x = n \mid S_0 = l) = \pi_l P(S_x = n \mid S_0 = l),$$

and

$$P(S_0 = n, S_x = l) = P(S_0 = n)P(S_x = l \mid S_0 = n) = \pi_n P(S_x = l \mid S_0 = n).$$

Since these two joint probabilities are equal,

$$\pi_l P(S_x = n \mid S_0 = l) = \pi_n P(S_x = l \mid S_0 = n).$$

Therefore,

$$P(S_x = l \mid S_0 = n) = \frac{\pi_l}{\pi_n} \left[1 - \rho_l(x \mid 0)\right]. \tag{16}$$

Since

$$\frac{\pi_l}{\pi_n} = \frac{\mu_l}{\mu_n},$$

we have

$$P(S_x = l \mid S_0 = n) = \frac{\mu_l}{\mu_n} \left[1 - \rho_l(x \mid 0)\right].$$

Therefore,

$$P(M_x \mid S_0 = n) = P_{\text{linker}} \frac{\mu_l}{\mu_n} \left[1 - \rho_l(x \mid 0)\right] + P_{\text{nuc}} \left[1 - \frac{\mu_l}{\mu_n} \left(1 - \rho_l(x \mid 0)\right)\right]. \tag{17}$$

Substituting Eqs. (14), (15), and (17) into Eq. (11) and simplifying gives

$$P(M_x \mid M_0) = A_l \left[\rho_l(x \mid 0) - \pi_l\right] + \rho_M, \tag{18}$$

where

$$A_l = \frac{\mu_l \left(P_{\text{linker}} - P_{\text{nuc}}\right)^2}{P_{\text{nuc}}\mu_n + P_{\text{linker}}\mu_l} \text{ and } \rho_M = \frac{P_{\text{nuc}}\mu_n + P_{\text{linker}}\mu_l}{\mu_n + \mu_l}. \tag{19}$$

Thus, deviations of the conditional methylation probability from the mean methylation level $\rho_M$ are proportional to deviations of the conditional linker-state probability from the mean linker probability $\pi_l$. At large distances,

$$\rho_l(x \mid 0) \to \pi_l,$$

and Eq. (18) therefore approaches the mean methylation probability $\rho_M$.

#### V. Histone-modification domain decay

The methylation model derived above assumes that positions 0 and $x$ lie within the same histone-modified region. We assume that GpCs outside histone-modified regions are unmethylated. Therefore, conditioning on a methylated GpC at position 0 implies that position 0 lies within a histone-modified region. For a second GpC at position $x$, the conditional methylation probability derived above applies only if position $x$ remains within the same histone-modified region as position 0.

This finite extent of histone-modified regions can produce a decay in the measured conditional methylation probability with increasing distance that is not captured by the nucleosome–linker model alone (Fig. S3). For example, for H3K27me3 at a CpG methylation fraction of 0.1–0.2, fitting the model without a domain boundary term ($\lambda = 0$) produces a systematic trend in the residuals with increasing distance (Fig. S3a). Allowing $\lambda$ to be fitted removes this trend. This effect is more pronounced for H3K4me3 at a CpG methylation fraction of 0–0.1, where the measured oscillations show a clear decay with distance that cannot be captured when $\lambda = 0$ (Fig. S3b). We therefore extend the

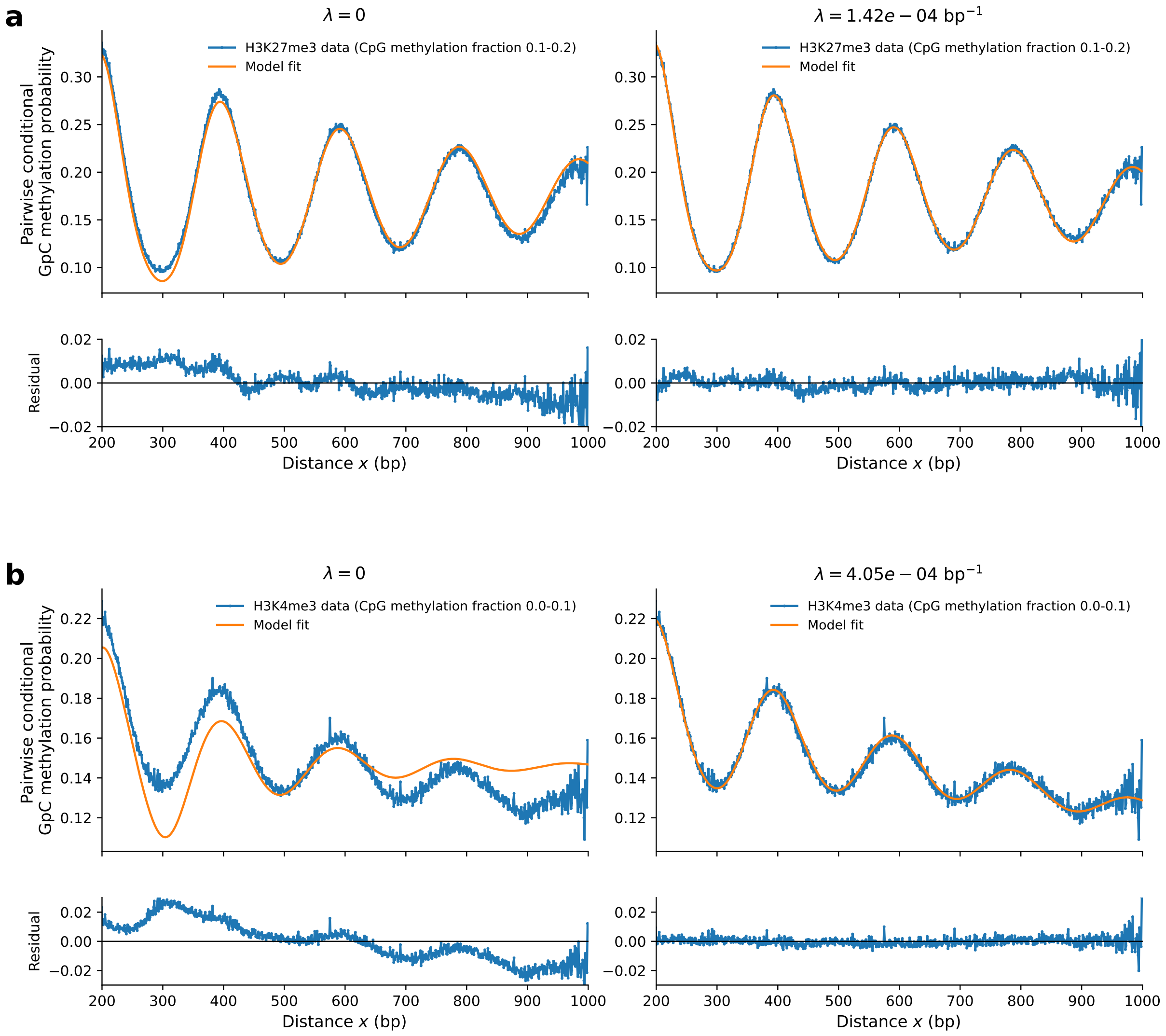


Figure S3. **Histone-modification domain boundaries capture the decay in methylation oscillations with distance. a.** Pairwise conditional GpC methylation probability for H3K27me3 at a CpG methylation fraction of 0.1–0.2, fitted with the nucleosome–linker model with $\lambda = 0$ (left) or with $\lambda$ fitted (right). Residuals between the measured probabilities and model fits are shown below. When $\lambda = 0$, the residuals show a systematic distance-dependent trend that is reduced when $\lambda$ is fitted. **b.** Corresponding comparison for H3K4me3 at a CpG methylation fraction of 0–0.1. In this example, the measured conditional methylation probability shows a pronounced decay with distance that is not captured by the model with $\lambda = 0$, whereas allowing $\lambda$ to vary captures this decay. Note that the residual y-axis limits differ between **a** and **b**.

model to account for the probability of leaving a histone-modified region as the distance from the methylated GpC at position 0 increases.

Our analysis is restricted to distances of at most 1 kb, corresponding to the length of the in silico fragmented reads. Therefore, we use a local approximation for the probability of encountering a histone-modified domain boundary over this distance. Specifically, we introduce a domain boundary probability $\lambda$, representing a constant probability per base pair of leaving the histone-modified region when moving from position 0 towards position $x$. Since the probability of encountering a boundary at any individual base pair is small, we approximate this probability as constant over the short genomic distances considered here. This local approximation does not require histone-modified domain lengths to follow an exponential distribution over longer genomic scales.

Under this local constant-boundary approximation, the probability of remaining within the same histone-modified region over a distance of $x$ base pairs is

$$P(D_x) = (1 - \lambda)^x,$$

where $D_x$ denotes the event that no domain boundary is encountered between positions 0 and $x$. For small $\lambda$,

$$(1-\lambda)^x = \exp\left[x\log(1-\lambda)\right] \approx \exp(-\lambda x).$$

We therefore approximate the probability of remaining within the same histone-modified region as

$$P(D_x) = e^{-\lambda x}. \tag{20}$$

GpCs outside the histone-modified region are assumed to be unmethylated. Therefore, the conditional methylation probability derived in Eq. (18) contributes only if no domain boundary is encountered between positions 0 and $x$. Multiplying Eq. (18) by the probability in Eq. (20) gives

$$P(M_x \mid M_0) = e^{-\lambda x}\left\{A_l\left[\rho_l(x\mid 0) - \pi_l\right] + \rho_M\right\}.$$

Substituting the expressions for $A_l$, $\pi_l$, and $\rho_M$ gives

$$P(M_x \mid M_0) = e^{-\lambda x}\left[\frac{\mu_l\left(P_{\text{linker}} - P_{\text{nuc}}\right)^2}{P_{\text{nuc}}\mu_n + P_{\text{linker}}\mu_l}\left(\rho_l(x\mid 0) - \frac{\mu_l}{\mu_n+\mu_l}\right) + \frac{P_{\text{nuc}}\mu_n + P_{\text{linker}}\mu_l}{\mu_n+\mu_l}\right]. \tag{21}$$

Together, Eq. (21) and the linker-state conditional probability in Eq. (10) define the analytical expression for the pairwise conditional methylation probability, which is fitted to the measured pairwise conditional methylation probabilities as described in Fig. 3 of the main text.

#### VI. Extension to two alleles

For fitting the single-cell oscillations, we extend the model to account for the presence of two alleles. The derivations above considers two GpCs located on the same DNA molecule. However, when measurements from the two alleles cannot be distinguished, a GpC at position $x$ may originate from either the same allele as the methylated GpC at position 0 or from the other allele. We assume that the two alleles contribute equally, such that each possibility occurs with probability 1/2. We denote the two alleles by subscripts (1) and (2). Without loss of generality, we let the methylated GpC at position 0 be on allele 1. The measured conditional methylation probability is thus

$$P(M_x \mid M_0) = \frac{1}{2}P(M_x^{(1)} \mid M_0^{(1)}) + \frac{1}{2}P(M_x^{(2)} \mid M_0^{(1)}). \tag{22}$$

Here, $P(M_x^{(1)} \mid M_0^{(1)})$ corresponds to the same-allele conditional methylation probability derived above. To calculate the second term, $P(M_x^{(2)} \mid M_0^{(1)})$ we introduce a parameter $k$ describing the concordance of the nucleosome–linker organization between the two alleles at the same genomic position,

$$k := P(S_0^{(2)} = l \mid S_0^{(1)} = l).$$

That is, $k$ is the probability that position 0 is in a linker state on allele 2 given that the same position on allele 1 is in a linker state. We assume that the linker length and nucleosome footprint distributions are described by the same parameters on both alleles. Furthermore, we assume that once the state at position 0 on allele 2 is specified, the state of allele 1 provides no additional information about the state at position $x$ on allele 2. Thus we have,

$$P(S_x^{(2)} = l \mid S_0^{(2)} = l) = \rho_l(x\mid 0),$$
$$P(S_x^{(2)} = l \mid S_0^{(2)} = n) = \frac{\pi_l}{\pi_n}\left[1 - \rho_l(x\mid 0)\right].$$

We first condition on the state of allele 2 at position 0

$$P(S_x^{(2)} = l \mid S_0^{(1)} = l) = k\,\rho_l(x\mid 0) + (1-k)\frac{\pi_l}{\pi_n}\left[1 - \rho_l(x\mid 0)\right].$$

Similarly,

$$P(S_x^{(2)} = l \mid S_0^{(1)} = n) = \beta\,\rho_l(x\mid 0) + (1-\beta)\frac{\pi_l}{\pi_n}\left[1 - \rho_l(x\mid 0)\right],$$

where $\beta = P(S_0^{(2)} = l \mid S_0^{(1)} = n)$. This $\beta$ can be expressed in terms of $\pi_l$, $\pi_n$, and $k$. Using the law of total probability,

$$\begin{aligned}\pi_l &= P(S_0^{(2)} = l)\\ &= P(S_0^{(2)} = l \mid S_0^{(1)} = l)\pi_l + P(S_0^{(2)} = l \mid S_0^{(1)} = n)\pi_n\\ &= k\pi_l + \beta\pi_n,\end{aligned}$$

and therefore

$$\beta = \frac{\pi_l}{\pi_n}(1-k).$$

We then define

$$c = \frac{k-\pi_l}{\pi_n}.$$

The conditional linker probabilities then become

$$\begin{aligned}P(S_x^{(2)} = l \mid S_0^{(1)} = l) &= \pi_l + c\left[\rho_l(x \mid 0) - \pi_l\right],\\ P(S_x^{(2)} = l \mid S_0^{(1)} = n) &= \pi_l - c\frac{\pi_l}{\pi_n}\left[\rho_l(x \mid 0) - \pi_l\right].\end{aligned}$$

As derived in previous sections,

$$\begin{aligned}P(S_0^{(1)} = l \mid M_0^{(1)}) &= \frac{P_{\text{linker}}\pi_l}{\rho_M},\\ P(S_0^{(1)} = n \mid M_0^{(1)}) &= \frac{P_{\text{nuc}}\pi_n}{\rho_M},\end{aligned}$$

where $\rho_M = \frac{P_{\text{nuc}}\mu_n + P_{\text{linker}}\mu_l}{\mu_n+\mu_l}$. Combining the two possible states of allele 1 at position 0 and using the above equations gives the conditional linker-state probability on allele 2

$$P(S_x^{(2)} = l \mid M_0^{(1)}) = \pi_l + c\frac{\pi_l(P_{\text{linker}} - P_{\text{nuc}})}{\rho_M}\left[\rho_l(x \mid 0) - \pi_l\right].$$

The methylation probability on allele 2 is

$$\begin{aligned}P(M_x^{(2)} \mid M_0^{(1)}) &= P_{\text{linker}}P(S_x^{(2)} = l \mid M_0^{(1)}) + P_{\text{nuc}}\left[1 - P(S_x^{(2)} = l \mid M_0^{(1)})\right]\\ &= \rho_M + c\frac{\pi_l\left(P_{\text{linker}} - P_{\text{nuc}}\right)^2}{\rho_M}\left[\rho_l(x \mid 0) - \pi_l\right]\\ &= \rho_M + c\frac{\mu_l\left(P_{\text{linker}} - P_{\text{nuc}}\right)^2}{P_{\text{nuc}}\mu_n + P_{\text{linker}}\mu_l}\left[\rho_l(x \mid 0) - \pi_l\right].\end{aligned}$$

The same-allele conditional probability derived in Eq. (18) is

$$P(M_x^{(1)} \mid M_0^{(1)}) = \rho_M + \frac{\mu_l\left(P_{\text{linker}} - P_{\text{nuc}}\right)^2}{P_{\text{nuc}}\mu_n + P_{\text{linker}}\mu_l}\left[\rho_l(x \mid 0) - \pi_l\right].$$

From Eq. (22) it then follows

$$P(M_x \mid M_0) = \rho_M + \frac{\mu_l\left(P_{\text{linker}} - P_{\text{nuc}}\right)^2}{P_{\text{nuc}}\mu_n + P_{\text{linker}}\mu_l}\frac{1}{2}\left[1 + \frac{k-\pi_l}{\pi_n}\right]\left[\rho_l(x \mid 0) - \pi_l\right].$$

Lastly, we assume that histone-modification domains are shared between the two alleles. Specifically, the same domain-boundary probability $\lambda$ applies to both the same and different allele contributions. Including the domain-boundary term gives

$$P(M_x \mid M_0) = e^{-\lambda x}\left[\frac{\mu_l\left(P_{\text{linker}} - P_{\text{nuc}}\right)^2}{P_{\text{nuc}}\mu_n + P_{\text{linker}}\mu_l}\frac{1}{2}\left[1 + \frac{\mu_n+\mu_l}{\mu_n}\left(k - \frac{\mu_l}{\mu_n+\mu_l}\right)\right] \times \left(\rho_l(x \mid 0) - \frac{\mu_l}{\mu_n+\mu_l}\right) + \frac{P_{\text{nuc}}\mu_n + P_{\text{linker}}\mu_l}{\mu_n+\mu_l}\right]. \tag{23}$$

When the nucleosome–linker states of the two alleles are independent, $k = \pi_l$, and the factor multiplying the oscillatory component $\left(\rho_l(x \mid 0) - \frac{\mu_l}{\mu_n+\mu_l}\right)$ becomes 1/2. In such case, the different allele pairs contain no information about the nucleosome state of the methylated anchor, which reduces the amplitude of the oscillatory signal by half. On the other hand, when the states of the two alleles are perfectly concordant, $k = 1$, this factor becomes one, which results in the single-allele model.

#### VII. Model fitting procedure

##### A. Fitting the model from the first oscillation peak

The nucleosome–linker model describes the periodic oscillations in pairwise conditional GpC methylation probability that emerge from nucleosome organization. However, the experimentally observed GpC methylation conditional probability curves at short genomic distances was not adequately captured by the model. Including these positions in the fitting procedure resulted in systematic deviations between the measured and fitted conditional methylation probabilities, as illustrated for H3K27me3 in Fig. S4a. We therefore fitted the model only from the first oscillation peak onwards, which greatly reduced the systematic deviations in residuals as shown in Fig. S4b. The first peak was identified from the measured pairwise conditional methylation probability, and positions preceding this peak were excluded from parameter inference. The deviations observed in the first ∼100bp might originate from proximity bias of the tethered methyltransferase which is not considered in the model.

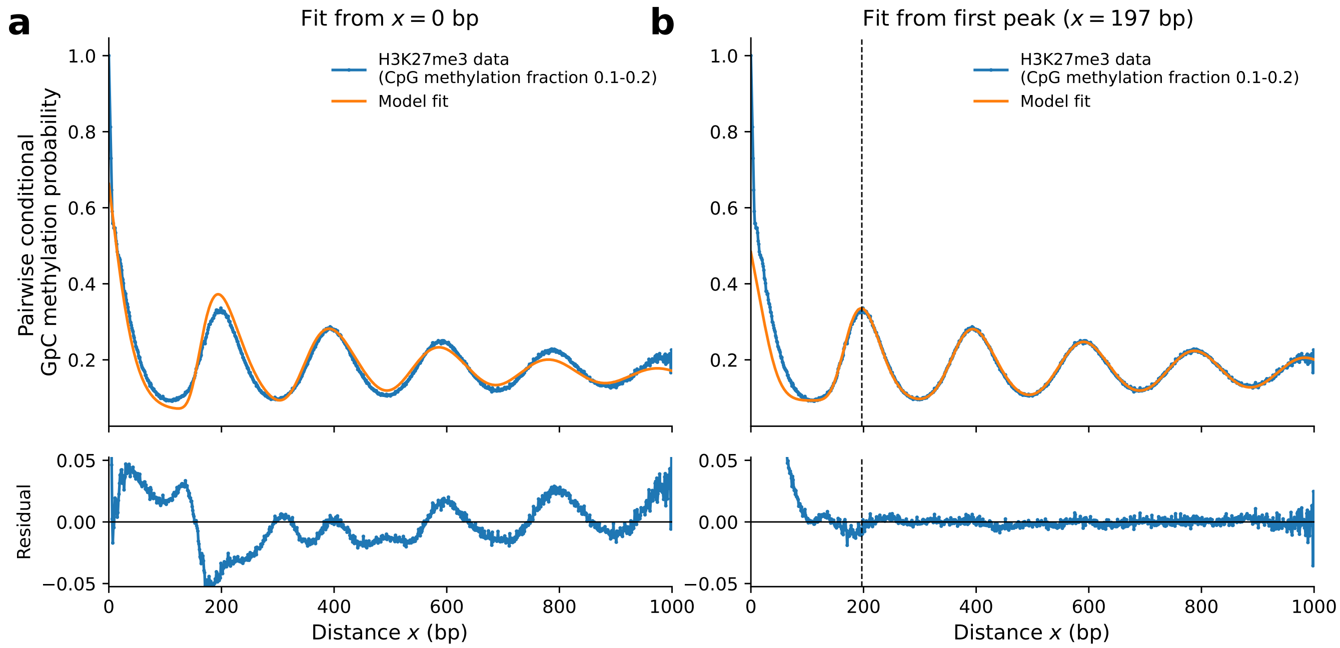


Figure S4. **The nucleosome–linker model does not accurately describe the observed data at short genomic distances.** Pairwise conditional GpC methylation probability for H3K27me3 at a CpG methylation fraction of 0.1–0.2, with residuals shown below. **a.** Model fitted over the complete distance range from $x = 0$ to 1000 bp. Inclusion of the short-distance region preceding the first oscillation peak results in systematic deviations between the measured and fitted probabilities. **b.** Model fitted from the first oscillation peak onwards. Positions preceding the first peak were excluded from parameter optimization but are shown for comparison. Restricting the fitting range reduces systematic deviations in the residuals over the fitted region.

##### B. Direct propagation of state probabilities

For numerical evaluation of Eq. (9), it is not necessary to explicitly construct the transition matrix $T$. Each substate has at at most two possible transitions, so multiplication by $T$ can instead be implemented directly using the probabilities $f_n(r)$ and $f_l(r)$.

Let $p_i(x)$ denote the entry of $\mathbf{p}_x$ corresponding to substate $i$, where nucleosome substates are indexed by $1, \ldots, N_n$ and linker substates by $N_n + 1, \ldots, N_n + N_l$. Propagation by one nucleotide is then given by

$$\begin{aligned}
p_{i+1}(x+1) &= p_i(x)\left[1 - f_n(i)\right], \qquad 1 \le i < N_n, \\
p_{N_n+1}(x+1) &= \sum_{i=1}^{N_n} p_i(x) f_n(i), \\
p_{i+1}(x+1) &= p_i(x)\left[1 - f_l(i - N_n)\right], \qquad N_n + 1 \le i < N_n + N_l, \\
p_1(x+1) &= \sum_{i=N_n+1}^{N_n+N_l} p_i(x) f_l(i - N_n).
\end{aligned}$$

Starting from $\mathbf{p}_0 = \mathbf{q}$, as defined in Eq. (8), these updates are applied successively for $x$ nucleotides. They are algebraically equivalent to the matrix propagation $\mathbf{p}_x^{\mathsf{T}} = \mathbf{q}^{\mathsf{T}} T^x$ in Eq. (9), but exploit the sparse structure of $T$ and avoid constructing the full transition matrix.

At each distance, the conditional linker-state probability in Eq. (10) is obtained directly by summing the probabilities of the linker substates,

$$\rho_l(x \mid 0) = \sum_{i=N_n+1}^{N_n+N_l} p_i(x).$$

##### C. Model fitting and parameter estimation

To determine the fitting interval, the experimentally measured pairwise conditional methylation probability was smoothed using a Savitzky–Golay filter with a window length of 50 points and a third-order polynomial. The first oscillation peak was identified as the maximum of the smoothed curve within indices 100–399, and only positions from this peak onwards were included in the fit, as described in Sec. VII A.

The mean nucleosome length was fixed at $\mu_n = 147$ bp. The remaining model parameters, $\mu_l$, $\sigma_n$, $\sigma_l$, $\lambda$, $P_{\text{linker}}$, and $P_{\text{nuc}}$, were fitted within the bounds

$$\begin{aligned}
20 &\le \mu_l \le 80 \text{ bp}, & 10 &\le \sigma_n \le 60 \text{ bp}, \\
10 &\le \sigma_l \le 60 \text{ bp}, & 0 &\le \lambda \le 0.001 \text{ bp}^{-1}, \\
0 &\le P_{\text{linker}} \le 1, & 0 &\le P_{\text{nuc}} \le 0.5.
\end{aligned}$$

For each candidate parameter set, the conditional state probabilities were calculated using the direct propagation described in Sec. VII B, and the corresponding pairwise conditional methylation probabilities were calculated from Eq. (21). Parameters were estimated by minimising

$$\text{RSS} = \sum_{x=x_{\text{peak}}}^{x_{\max}} \left[P_{\text{model}}(M_x \mid M_0) - P_{\text{measured}}(M_x \mid M_0)\right]^2$$

using differential evolution. Independent optimisation rounds were performed for each histone modification and CpG methylation fraction, and the solution with the lowest RSS was retained.

Parameter uncertainties were estimated locally around the optimal solution using a numerical Jacobian calculated by centred finite differences. The covariance matrix of the free parameters was approximated as

$$\mathbf{C} = s^2 \left(\mathbf{J}^{\mathsf{T}}\mathbf{J}\right)^+, \qquad s^2 = \frac{\text{RSS}}{N - p},$$

where $\mathbf{J}$ is the Jacobian evaluated over the fitted distance range, $(\cdot)^+$ denotes the Moore–Penrose pseudoinverse, $N$ is the number of fitted distance points, and $p$ is the number of free parameters.
